# Early Detection and Quantification of Fusarium Wilt in Greenhouse-Grown Tomato Plants Using Water-Relation Measurements

**DOI:** 10.1101/2024.02.11.579801

**Authors:** Shani Friedman, Ahan Dalal, Dor Batat, Saul Burdman, Yheonatan Sela, Matanel Hipsch, Shilo Rosenwasser, Evgeniya Marcos Hadad, Shay Covo, Menachem Moshelion

## Abstract

Visual estimates of plant symptoms are traditionally used to quantify disease severity. Yet, the methodologies used to assess these phenotypes are often subjective and do not allow tracking of disease progression from very early stages. Here, we hypothesized that quantitative analysis of whole-plant physiological vital functions can be used to objectively determine plant health, providing a more sensitive way to detect disease. We studied the tomato wilt that is caused by *Fusarium oxysporum* f. sp. *lycopersici*. Physiological performance of infected and non-infected tomato plants was compared using a whole-plant pot-based lysimeter functional-phenotyping system in a semi-environmentally controlled greenhouse. Water-balance traits of the plants were measured continuously and simultaneously in a quantitative manner. Infected plants exhibited early reductions in transpiration and biomass gain, which preceded visual disease symptoms. These changes in transpiration proved to be effective quantitative indicators for assessing both plant susceptibility to infection and virulence of the fungus. Physiological changes linked to fungal outgrowth and toxin release contributed to reduced hydraulic conductance during initial infection stages. The functional-phenotyping method objectively captures early-stage disease progression, advancing plant disease research and management. This approach emphasizes the potential of quantitative whole-plant physiological analysis over traditional visual estimates for understanding and detecting plant diseases.

## Introduction

Pathogens cause severe losses to agricultural production. Worldwide, an estimated 20 to 40% of crop yields are lost to pests and plant diseases (CABI n.d.). Each year, plant diseases cost the global economy around $220 billion (Sarkozi 2019). Therefore, monitoring plant health and detecting pathogens at an early stage are crucial for reducing pathogen establishment and spread, and for facilitating effective management practices in sustainable agriculture (Martinelli et al. 2015).

While in agricultural settings, disease severity is determined by yield or economic loss; in plant pathology research, the definition is more complex. Disease severity is the proportion of the experimental plant unit exhibiting visible disease symptoms and is often expressed as a percentage (Madden et al. 2007). The visual estimations that are often used to quantify disease severity are subjective and prone to inaccuracy that can lead to incorrect conclusions (Bock et al. 2020; Stewart and McDonald 2014). Objective quantitative assessment of plant disease severity is critical for accurately determining pesticide efficacy, correlating yield losses with disease damage, calculating damage thresholds, and conducting reproducible experiments in any plant–pathogen interaction study. A quantitative method for studying disease progression would help us to understand the contribution of each determinant (plant, pathogen, and environment) to the complex phenotype of a plant disease.

Existing spectroscopic and image algorithm-based disease diagnostic techniques, such as RGB imaging, multispectral and hyperspectral sensors, thermography, and chlorophyll fluorescence, rely on standardized protocols and complex data analysis methods, which can be a barrier for practical application. Additionally, the interpretation of imaging data often requires expert knowledge to ensure accuracy and reliability (Mahlein 2016). Gaunt (1995) previously suggested that adopting a simpler way to measure disease, based on deviations from normal plant function, could potentially resolve many of the challenges faced in linking disease with yield (Gaunt 1995).

Healthy plants growing under optimal conditions adapt to environmental variations in ways that optimize their productivity. For instance, a healthy plant adeptly adjusts its stomatal opening to maximize CO_2_ absorption when it is exposed to favorable ambient conditions, such as sufficient light, adequate soil moisture, and optimal temperature (Chaves et al. 2003; Schroeder et al. 2001). In contrast, a diseased plant will respond differently under identical conditions, potentially failing to exploit the environment to maximize its productivity potential (Jones 2013). Plant-pathogenic fungi can decrease hydraulic conductance or transpiration in plants through various mechanisms. One mechanism is the disruption of water uptake and transport associated with the clogging of the xylem vessels by fungal mycelia (reviewed by Nemec et al. 1986; Srinivas et al. 2019). Another mechanism is the secretion of polysaccharides and toxins by the fungus into the xylem, which are subsequently transported to the leaves, and have been shown to reduce transpiration and induce other physiological perturbations (Singh et al. 2017). Specific substances contributing to these effects include chitin (Attia et al. 2020) and fusaric acid (Dong et al. 2012).

In this study, we focused primarily on the *Fusarium oxysporum* f. sp*. lycopersici* - tomato pathosystem. *F. oxysporum* f. sp*. lycopersici* is a soil-borne fungus that spreads in the tomato plant through the vascular tissues, causing the plant to wilt. This phenomenon is assumed to be associated with both physical clogging of the xylem vessels and toxins secreted by the fungus into the xylem (Kashyap et al. 2021). Physical and chemical barriers contribute to vascular blockage, disrupting plant water-balance and promoting wilting (Kashyap et al. 2021). Given the significant impact of *F. oxysporum* f. sp*. lycopersici* on water-related physiological traits, we hypothesized that these traits might serve as early markers of infection in the leaf and the whole plant. We hypothesize that careful examination of subtle physiological changes (i.e., suboptimal functioning of the plant) that occur early in the infection process can provide insights into pathogen infection before the later emergence of visible symptoms. Furthermore, we hypothesize that suboptimal physiological parameters, such as reduced transpiration, can signal dysfunction resulting from the interaction with the pathogen in a more quantitative manner than visual assessment alone.

Our method focuses on continuous direct monitoring of critical physiological parameters, such as transpiration rates and biomass changes, which are easy to use and interpret. However, this approach has not been widely used for early detection of disease (Bock et al. 2020; Fang and Ramasamy 2015; Martinelli et al. 2015). Some studies have focused on the assessment of water-balance parameters to quantify and compare disease severity (Dong et al. 2012; Wang et al. 2015); however, these studies involve laborious, less-precise and time-consuming techniques.

The objectives of this study were: (i) to develop and validate a high-throughput physiological phenotyping approach for early disease detection in plants under semi-controlled greenhouse conditions; (ii) to test the sensitivity of physiological traits, such as transpiration rates and biomass changes, for quantifying *F. oxysporum* f. sp. *lycopersici* virulence and tomato susceptibility; (iii) to investigate how *F. oxysporum* f. sp. *lycopersici* impacts transpiration, by testing the impact of its toxins on leaf hydraulic conductance; and (iv) to test the generalizability of this approach using an additional pathosystem, potato (*Solanum tuberosum*)-*Phytophthora infestans* Mont. DeBary (the late blight disease). This work might contribute to a broader understanding of plant–pathogen–environment interactions and help to improve the methods available for studying and early detection of vascular wilt diseases.

## Materials and Methods

### Greenhouse conditions

This work was performed in a semi-controlled greenhouse located at the Robert H. Smith Faculty of Agriculture, Food and Environment, Hebrew University (Rehovot, Israel), which is an extension of the I-CORE Center for Functional Phenotyping greenhouse (http://departments.agri.huji.ac.il/plantscience/icore.phpon) for plant disease research. The greenhouse allows natural day length (10 to 14 hours daylight) and light conditions (photosynthetic photon flux density of 500 to 1000 μmol/m^2^/s^1^ at mid-day) and has a desert cooler along its northern wall to prevent overheating. During summer, the average temperature in the greenhouse is 32.86°C during the day and 27.35°C at night, with relative humidity averaging 59.20% and 79.93%, respectively. During the winter, the daytime temperature averages 22.10°C, dropping to 13.48°C at night, with corresponding relative humidity levels of 50.46% and 77.04%. Environmental data, such as PAR light, temperature, relative humidity and vapor pressure deficit, were monitored continuously and recorded every 3 min (Fig. S1).

### Whole-plant physiological phenotyping measurements

Whole-plant, continuous physiological measurements were taken using load cells (lysimeters) (PlantArray 3.0 system; Plant-DiTech, Yavne, Israel). In the PlantArray system each pot is placed on a gravimetric load cell, which is connected to a control unit that continuously measures the pot mass. This setup allows for precise calculation of various physiological traits. (For more information, please see Fig. 1 and Dalal et al. 2020; Halperin et al. 2017). In the greenhouse, there were 36 to 60 PlantArray units, which collected data and controlled the irrigation for each pot separately. The data were analyzed using SPAC-analytics (https://spac.plant-ditech.com), a web-based software program that allows analysis of the real-time data collected from the PlantArray system (Halperin et al. 2017).

**Fig. 1.**
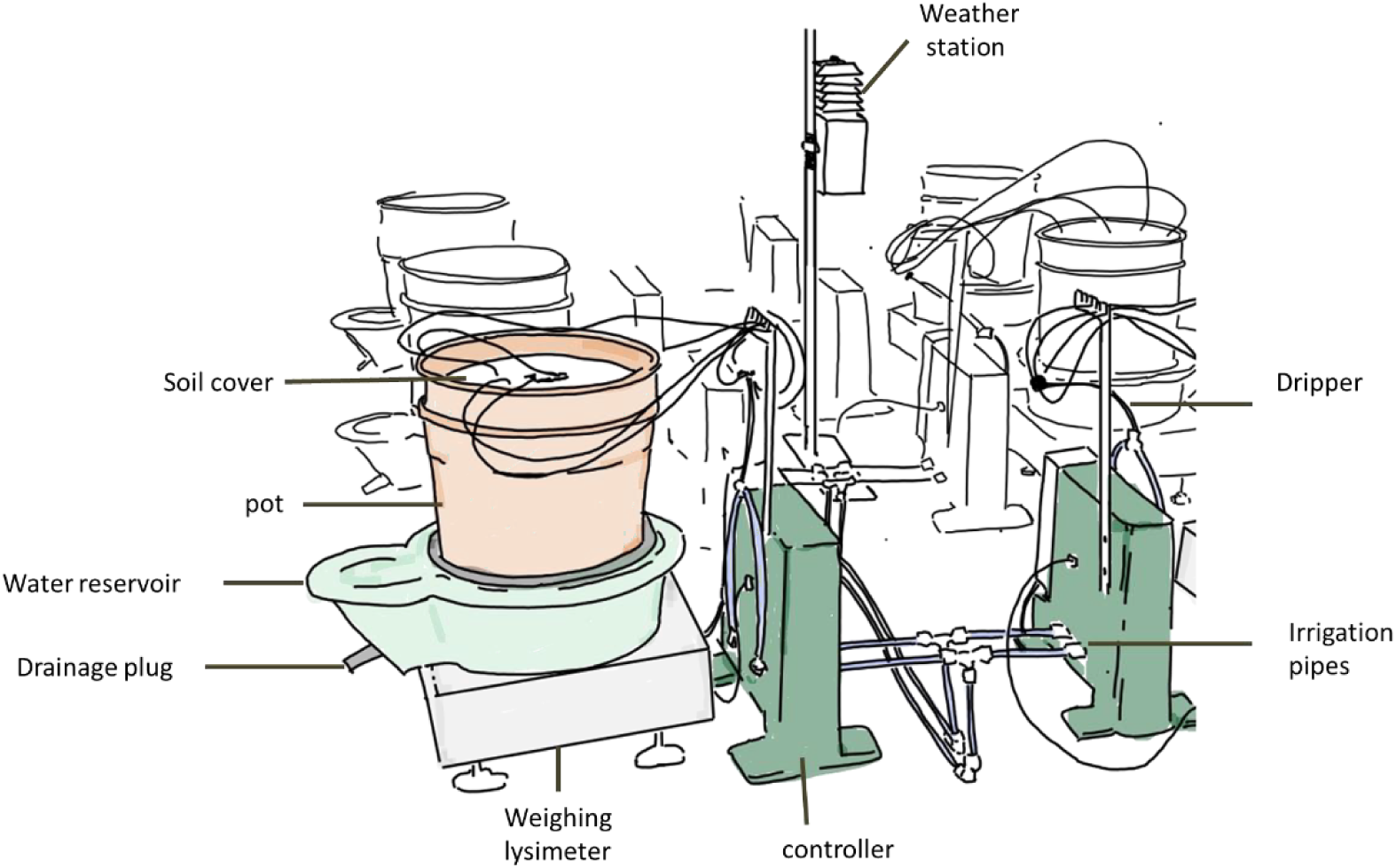
PlantArray System Description. The PlantArray system is a telemetric gravimetric platform designed for real-time physiological phenotyping of plants. It includes multiple weighing lysimeters and environmental sensors for non-destructive, high-throughput measurements of soil, plant, and atmospheric conditions, as detailed by Dalal (2021) and presented here briefly. Each plant is placed in a pot within a green container that collects drainage water. A white cover and a black rubber ring minimize evaporation, ensuring water loss occurs primarily through transpiration. The pots are situated on weighing lysimeters, where the lysimeter’s load cell converts mass into an electrical charge. A metal platform ensures accurate measurement. The controller unit regulates irrigation and connects to load cells and sensors, while metal stands support pipes and cables to avoid interference with measurements. The weather station’s atmospheric sensors measure relative humidity, temperature, vapor pressure deficit, and photosynthetically active radiation. These measurements are crucial for assessing environmental conditions and plant physiology, such as stomatal conductance.

Pots, sand, and plants were weighed ahead of the experiment. The pots were covered with a custom cover that only allowed the stem through it, to minimize evaporation. Plants were irrigated nightly between 22:00 and 02:00 using a drip irrigation system that delivers water to plant root zones via a controlled network of valves and emitters. The system performs multiple irrigation cycles followed by drainage periods to consistently return the substrate to field capacity (referred as to well-irrigated in this article). This multicycle watering and drainage ensures optimal soil moisture conditions and promotes proper leaching to remove excess salts. Daily pre-dawn pot mass was measured after full drainage. Plant mass was calculated at this point. The difference in pot mass between consecutive days reflects the plant’s biomass gain. The lack of additional irrigation throughout the daylight hours ensured a monotonic pot-mass decrease between subsequent irrigation events.

Transpiration, referred to as ‘daily transpiration’ or ‘transpiration rate’ and E (calculated in terms of water transpired per unit of plant net mass per min), were calculated based on the mass loss during the day. For full the details on calculations and system see Dalal et al. (2020) and Halperin et al. (2017).

### Tomato plant material

To assess whether physiological traits quantitatively correlate with tomato cultivar susceptibility to *F. oxysporum* f. sp*. lycopersici* race 2, we selected several tomato cultivars previously categorized on a susceptibility-tolerance-resistance scale in response to this pathogen. These included the commercial cultivar *Lycopersicon esculentum* M82 (Eshed and Zamir 1995), described as tolerant to this pathogen (Sela-Buurlage et al. 2001) and the cultivars Rehovot-13 (Hazera Genetics, Brurim Farm, Israel) and Marmande Verte, which are considered susceptible. In addition, we used the nearly isogenic tomato cultivars Moneymaker and Motelle, referred to as susceptible and resistance to *F. oxysporum* f. sp*. lycopersici* race 2, respectively (Sarfatti et al. 1989).

### *F. oxysporum* f. sp*. lycopersici* strains and growth conditions

To evaluate the ability of physiological traits to quantify fungi virulence, we selected two isolates of *F. oxysporum* f. sp*. lycopersici* race 2: *F. oxysporum* f. sp*. lycopersici* 4287 obtained from the American Type Culture Collection (ATCC) and fr2T obtained from Hazera Genetics. In preliminary experiments, *F. oxysporum* f. sp. *lycopersici* 4287 was found to be a moderately virulent strain (mvF) and fr2T was found to be a highly virulent strain (vF). The fungi were grown on plates containing 39 g/liter potato dextrose agar (PDA; BD Difco, NJ, USA) with additional 5 g/liter Bacto Agar (Difco, MD, USA). For formation of microconidia, the strains were grown in a liquid minimal medium (LMM) that contained 100 mM KNO3, 3% sucrose and 0.17% yeast nitrogen base without amino acids (Cohen et al. 2019). *F. oxysporum* f. sp*. lycopersici* was cultured in 500 ml Erlenmeyer flasks containing 50 ml LMM. The inoculum was obtained either from PDA plates (1 cm² plugs) or from stored aliquots (50 µl). The Erlenmeyer flasks were incubated for 6 days at 28°C in an incubator (New Brunswick Innova42/42R, Hamburg, Germany) without internal lighting or light shielding, with rotary shaking at 185 rpm. The resulting fungal cultures were filtered through a 40 µm sterile strainer (Corning, Cell Strainer; Sigma-Aldrich, Rehovot, Israel) to remove the mycelia and then centrifuged at 3,220 g for 10 min to pellet the conidia. The conidia were resuspended in sterile water and quantified using a Neubauer hemocytometer counting chamber (Mariefeld, Lauda-Königshofen, Germany) under a Nikon H550S microscope (Nikon, Tokyo, Japan). *F. oxysporum* f. sp. *lycopersici* was stored for long periods at −80°C, in aliquots containing 30% glycerol and about 10^9^ cells/ml.

### Greenhouse experimental set up

Tomato seeds were germinated in 50 ml cavity trays (Histhil, Nehalim, Israel) filled with soil (Matza Gan; Shacham, Givat-Ada, Israel) in the greenhouse. They were fertigated with ‘Mor’ (CIL, Beer Sheva, Israel) water every 2 to 3 days until germinated and then irrigated daily until transferring to the bigger pots. Three- to 4-week-old tomato plants were inoculated; the plant roots were washed with tap water and trimmed to leave 5 cm of root length, ensuring a more uniform and homogeneous infection. The roots were then submerged in a suspension containing *F. oxysporum* f. sp. *lycopersici* (10^7^–10^6^ conidial spores/ml distilled water) for 5 to 10 min. The plants were then transplanted into pots at pot water capacity. Control plants were treated similarly, except that sterile water was used instead of the conidial suspension.

The plants were then transplanted into 4 liter pots filled with Silica sand (grade 20-30, particle size 0.595–0.841 mm; Negev Industrial Minerals Ltd., Yeruham, Israel). In one experiment, plants were grown in a commercial growth medium (Matza Gan; Shacham) to assess its potential impact on symptom development (Experiment 4, Tables 1 and S1). On the day of transplanting, the soil was kept at field capacity to ensure the plants’ establishment. From the following day, irrigation was applied only at night. The irrigation was with fertilized water (‘Mor’; CIL). The pots were set on the PlantArray system and were monitored continuously for about 30 to 40 days.

The pots were arranged on tables, with each table containing 12 pots. Treatments were assigned using a random block design within each table to minimize spatial effects and ensure unbiased distribution of treatments across the greenhouse. Each treatment group was replicated four to six times across the experiment to ensure statistical use (Fig. S2).

### Visual disease severity scoring

Disease severity was monitored continuously (daily) and was scored with the following symptom severity scale (Fig. S3): 0, asymptomatic plants; 1, weakly symptomatic plants (<25% of leaves chlorotic or wilted); 2, moderately symptomatic plants (25 to 50% of leaves were chlorotic or wilted); 3, highly symptomatic plants (>50% of the leaves wilted, but plants were alive); and 4, dead plants. The area under the disease progress curve (AUDPC) per pot was calculated. At the end of the experiment, 3 to 4 weeks after inoculation, we measured both the fresh weight and the height of the plant shoots and, in some cases, the roots as well.

### Pathogen-progress stem sampling assays

At the end of every experiment to validate *F. oxysporum* f. sp. *lycopersici* infection, we conducted a series of post-harvest analyses. These included cutting the plant at the stem base to assess the browning of the vascular system, a typical symptom induced by *F. oxysporum* f. sp. *lycopersici* in tomato (Rep et al. 2005). Next, we performed fungal outgrowth tests by collecting stem pieces from each plant at 0, 5, and 15 cm above the soil surface to confirm that the observed symptoms were caused by the applied treatments. These stem pieces were then surface-disinfested by briefly submerging them in 70% ethanol and igniting the ethanol by passing the stems through an open flame. A slice of each sterilized stem piece was put on a Petri dish (90 mm) filled with PDA supplemented with 250 mg/liter streptomycin or 100 µg/ml ampicillin to reduce bacterial contamination (van der Does et al. 2019); both antibiotics exhibit similar inhibitory activity. After 4 days of incubation in the dark at 28°C, fungal outgrowth was assessed using a double-blind method, in which the persons examining the samples did not know what the treatments were thus ensuring a non-biased evaluation. The assessment focused on comparing fungal growth texture, color, and overall morphology, with mycelia of *F. oxysporum* f. sp. *lycopersici* being characterized by a fluffy texture and a pink/violet tan coloration (Fig. 2b).

**Fig. 2.**
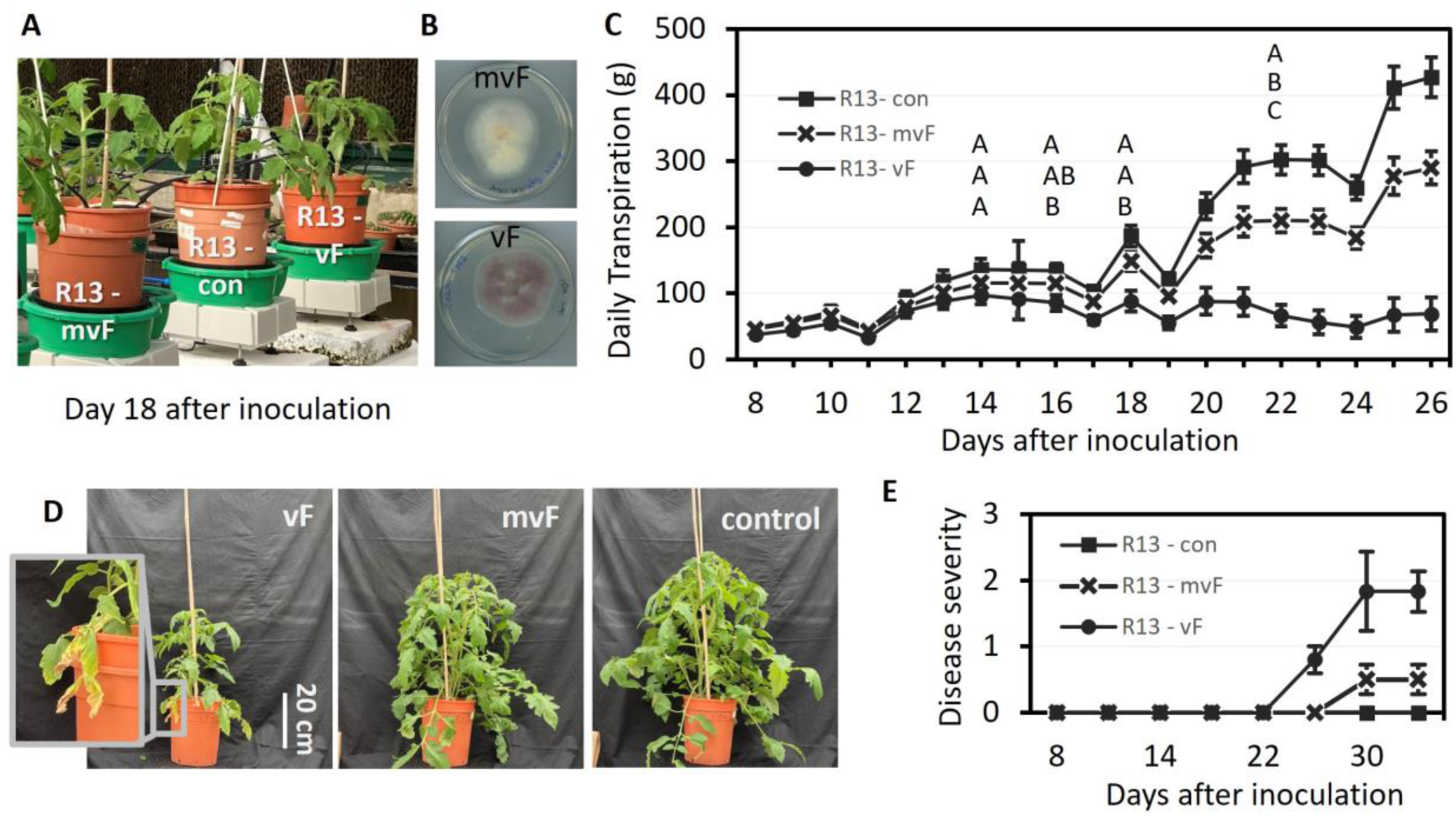
Variations in transpiration corresponded to virulence levels prior to visible signs of disease. Rehovot-13 (R13) susceptible plants infected with either mvF (common strain) or with vF (virulent strain) were compared with control (mock-infected plants). A, Plants at day 18 after inoculation in the PlantArray system. B, At the end of the experiment, stem pieces were incubated on PDA plates. The outgrowth of fungi was transferred to new plates; mvF fungi were light pink while vF fungi were violet. All infected plants showed stem browning and fungal outgrowth, while none of the controls did. C, Daily transpiration of the whole plants (mean ± SE), different letters indicate a significant difference between treatments; *P* ≤ 0.05. Differences were tested, and letters were assigned only in the first instance when significant changes occurred. D, Plants at the end of the experiment (Day 39 after inoculation). E, Disease progress curve, error bars indicate standard error. Average of wilt symptoms evaluated using the disease-severity index described in Materials and Methods. *N* = 6. The data presented here are from experiment number 2 as listed in Table 1.

**Table 1.** Summary of F. oxysporum f. sp. lycopersici –tomato experiments: Plant transpiration, mass, and disease symptoms.

| Experiment |  | Treatment |  | Symptoms |  |  | Duration of exp. |
| --- | --- | --- | --- | --- | --- | --- | --- |
| Exp. | Months | *Fusarium* | Tomato | Transpiration<sup>a</sup> | Mass<sup>a</sup> | Morphology<sup>b</sup> |  |
| 1A | Oct-Nov | vF | M82 | 18 days | 31 days | - | 39 days |
| 2 | Dec | vF | M82 | 24 days | 35 days | - | 39 days |
| 3 | Apr | vF | MM | 13 days | 17 days | - | 19 days |
| 1B | Oct-Nov | mvF | R13 | 6 days | 19 days | 21 days | 75 days |
| 4 | Aug | mvF | R13 | 11 days | 25 days | 60 days | 34 days |
| 2 | Dec | mvF | R13 | 22 days | 32 days | 32 days | 39 days |
| 2 | Dec | vF | R13 | 16 days | 25 days | 28 days | 39 days |
| 5 | Jan | vF | MV | 22 days | 7 days | 35 days | 42 days |

The plants tested included M82 (tolerant), R13 (susceptible), Mv (susceptible) and MM (susceptible). Plants were inoculated with either the moderately virulent strain (mvF) or the highly virulent strain (vF).
<sup>a</sup> “Transpiration” and “mass” columns indicate the point in time (dai) at which a t-test revealed a significant difference from the control; $p < 0.05$ .
<sup>b</sup> “Morphology” column indicates the point in time (dai) at which 50% of plants exhibited chlorosis or wilting.

### Molecular confirmation of fungal identity

Verification of pathogen occurrence in infected plant tissue was carried out by PCR targeting *F. oxysporum* f. sp. *lycopersici*-sequences. Fungal plugs (1cm^2^) from the stem outgrowth assay (see above) were grown for 5 days in PDA at 28°C under static conditions. Mycelia were harvested and ground with a pestle and mortar with liquid nitrogen. Samples were stored at -20°C overnight. Genomic DNA was extracted using either the Hi PurATM Fungal DNA Mini Kit (HiMedia, Maharashtra, India), following the manufacturer’s instructions, or the CTAB method (Muraguchi et al. 2003; Zolan and Pukkila 1986).

The DNA samples were stored at −20°C until further use. Samples of genomic DNA (3 µl) were added to PCR tubes containing appropriate primers (as described below), PCR-grade H_2_O and RedTaq Ready Mix (R2523, Sigma-Aldrich, Rehovot, Israel) according to the manufacturer׳s instructions. The amplified products were subjected to electrophoresis on 1.2% agarose gel containing agarose in 0.5 X TBE buffer and were stained with ethidium bromide. Gel images were captured using a C200 gel imaging workstation (Azure Biosystems; Dublin, CA, USA).

The amplification protocol used for the PCR reactions included an initial denaturation at 94°C for 1 min, followed by 25 cycles of denaturation at 94°C for 1 min, annealing at 58°C for 30 s, extension at 72°C for 1 min, and a final extension at 72°C for 7 min. The target genes amplified were the *six5* gene for *F. oxysporum* f. sp. *lycopersici* (product size: 667 bp) and the nrDNA ITS region (product size: 340 bp). The *six5* gene primers were *six5*-f1 (5’-ACACGCTCTACTACTCTTCA-3’) and *six5*-r1 (5’-GAAAACCTCAACGCGGCAAA-3’) (Taylor et al. 2016). The primers for the ITS region were FOF (5’-ACATACCACTTGTTGCCTCG -’) and FOR (5’- CGCCAATCAATTTGAGGAACG-3’) (Prashant et al. 2003).

### Seedling assays for virulence testing

To test the quantitative ability of the system to assess *F. oxysporum* f. sp. *lycopersici* virulence, we initially performed a standard pathogenicity assay on the two isolates used in this study: *F. oxysporum* f. sp. *lycopersici* 4287 (mvF) and fr2T (vF). Tomato seeds were sown in vermiculite V2U (Agrekal, Moshav Habonim, Israel) and seedlings were grown for 10 to 13 days in a temperature-controlled glasshouse with a maximum daytime temperature of 25°C and minimum nighttime temperature of 18°C prior to fungal inoculation. *F. oxysporum* f. sp. *lycopersici* cultures were grown and filtered as described above. Seedlings were removed and their roots were washed with water, trimmed, and then dipped in a conidial suspension (16 × 10^6^ conidia/mL) for 5 min before being replanted in a mixture of vermiculite with autoclaved soil (4 units vermiculite:1 unit soil). Roots of control plants were dipped in water for the purposes of mock inoculation, to ensure that disease phenotypes were a consequence of *F. oxysporum* f. sp. *lycopersici* infection rather than the inoculation process per se. Plants were grown in trays; each tray contained both susceptible plants (cv. Moneymaker) and resistant (cv. Motelle) controls to ensure infection. To ensure successful infection and development of disease symptoms, the seedlings were densely planted and covered with foil for 2 days, with no fertilizer applied. Plants were kept in a controlled-environment growth room at 27°C with a 16/8 h day/night cycle.

After 18 or 21 days, wilt symptoms and vascular browning were recorded and used to calculate disease scores according to criteria similar to those described by Rep et al. (2005): 0, healthy plant; 1, one or two brown vascular bundles in the hypocotyl; 2, at least two brown vascular bundles and growth distortion; 3, all vascular bundles are brown; plant either dead or very small and wilted (Supplementary Fig. S4).

### Impact of *F. oxysporum* f. sp. *lycopersici* filtrate (toxins) on leaf hydraulic conductance (K_leaf_)

To investigate how *F.oxysporum* f. sp. *lycopersici* impacts transpiration, we tested the impact of the fungal toxin extract on leaf hydraulic conductance. Fungal culture filtrate was perfused into the leaf vascular system and compared to positive controls, including chitin and abscisic acid (ABA), which are known to reduce K_leaf_ (Attia et al. 2020).

To prepare the *F. oxysporum* f. sp*. lycopersici* filtrate toxin, the strains were cultured in Czapek Dox broth, as described in previous studies (Portal et al. 2018; Scala et al. 1985; Sutherlandt and Pegg 1992). Briefly, 50 ml of Czapek Dox broth was inoculated with (for culture filtrate) or without (for control filtrate) the strains, and then incubated on an orbital shaker set at 130 rpm in darkness for 14 days at 25°C. The cultures were then filtered through Whatman filter paper and the filtrate concentrated up to approximately 10% of the initial volume by rotary evaporation at 25°C. The concentrated filtrates were filtered through 0.45μm sterile syringe filters, to remove any spores or mycelial fragments (for culture filtrate) that might cause physical blockage during the petiole-fed treatments for the hydraulic experiment.

Tomato plant leaflets, approximately 1.5 to 2 months old, were selected from the first three leaflets from top younger leaves of similar size. Only leaflets with no noticeable injuries or anomalies were used. Before dawn, these leaflets were cut and immediately placed in a 2 ml tube (Eppendorf, Hamburg, Germany), with their petioles dipped in artificial xylem solution (AXS; Attia et al. 2020). The AXS was supplemented with one of the following treatments: 10 μM abscisic acid (ABA), 0.2 mg/ml chitin prepared from a 10 mg/ml stock solution according to Attia et al. (2020), 0.5% of the culture filtrate, or a control filtrate. The leaflets were exposed to a light intensity of 150 µmol/m^2^ s^2^ at 26 to 28°C. Transpiration from each perfused leaf and its water potential were measured between 2 to 4 h of treatment using an Li-600 porometer (LI-COR, Bnei Brak, Israel) and pressure chamber (Arimad-3000; MRC Ltd., Holon, Israel), respectively, as described previously (Grunwald et al. 2021). Leaf hydraulic conductance (K_leaf_) was calculated as the negative of the ratio of transpiration to water potential (Grunwald et al. 2021).

### Testing the method generalizability using the potato–*Phytophthora infestans* pathosystem

To test the generalizability of physiological parameters as sensitive indicators of disease in other plant-pathogen systems, we evaluated their applicability using *Solanum tuberosum* cv. *Desiree* (Hipsch et al. 2021) and *Phytophthora infestans* isolate 164 (genotype 23_A1, resistant to mefenoxam). This isolated was collected in March 2016 from a potato field in Nirim, in the western Negev, Israel (Cohen 2020). The isolate was propagated in a growth chamber at 18°C by repeated inoculations of freshly detached potato leaves (Hipsch et al. 2023). For inoculation, freshly produced sporangia were collected at 5 to 7 days after inoculation (dai) from infected leaves by washing with distilled water (DW) into a beaker kept on ice (4°C).

Potato plants were vegetatively propagated from cuttings or tubers and placed in moist soil (Green 761, Even Ari, Beit Elazari, Israel) in 26.82 × 53.49 cm pots within a controlled environment greenhouse. For the experiment, 3- to 4-week-old plants were transferred from the controlled-environment greenhouse to the I-CORE greenhouse facility, where they were placed on the PlantArray system in 4-liter pots filled with silica sand (grade 20–30; Negev Industrial Minerals Ltd.). We inoculated the potato plants by spraying their foliage with a fresh suspension of *P. infestans* (genotype 23_A1) sporangia (1 × 10^5^ sporangia/ml). The plants were then covered with black plastic bags and incubated in the dark for 18 h until the next morning, then the bags were removed (Hipsch et al. 2023). Plants sprayed with DW served as the control. This experiment was conducted once in the greenhouse during winter. On the day of inoculation, the temperature reached a maximum of 24°C at noon and dropped to a minimum of 10°C at night.

### Statistical analyses

We used the JMP® ver. 16 statistical packages (SAS Institute, Cary, NC, USA) for our statistical analyses. Student’s t-test was used to compare the means of two groups and ANOVA was used to compare means among three or more groups. The greenhouse experiment treatments were randomly distributed to account for the potential effects of plant placement on the results (as illustrated in Fig. S2). The main effects analyzed included treatment type (e.g., fungal strain or control) and plant genotype (resistant or susceptible cultivars). The homogeneity of variances across treatments was tested using Levene’s test to ensure the validity of ANOVA assumptions. Differences between the treatments were examined using Tukey’s HSD test. Each analysis involved a set significance level of *P* < 0.05

## Results

### High-throughput whole plant physiological measurement sensitivity for early detection of *F. oxysporum* f. sp. *lycopersici* infection

This research focused on the use of the PlantArray system to detect and quantify early effects of disease in plant–pathogen interactions. The system monitors real-time physiological responses, including transpiration and biomass changes (see Methods for details). This study primarily involved two strains of *Fusarium oxysporum* f. sp. *lycopersici*: a moderately virulent strain (mvF) and a highly virulent strain (vF).

To test the sensitivity of the PlantArray system we conducted several independent repetitions with these strains, inoculating different tomato cultivars (Table S1, Table 1). Experiments resulting in any type of symptom, physiological or morphological, are presented in Table 1. All experiments that were conducted in our semi-controlled greenhouse are presented in Table S1. In 11 out of 19 experiments (Table S1), no visible symptoms or alterations in transpiration were observed, despite successful infection that was confirmed by fungal outgrowth tests. In seven out of eight experiments in which symptoms were observed (reduction in daily transpiration, reduction in plant mass or visual symptoms), the infection was first detected by a reduction in daily transpiration, followed by other parameters (Table 1).

The first evidence of disease in Rehovot-13 (R13) plants inoculated with *F. oxysporum* f. sp. *lycopersici* was consistently a reduction in daily transpiration (6 to 22 dai), followed by a decrease in plant mass (19 to 32 dai) and, eventually, visible symptoms (21–60 dai; Table 1). No visible disease symptoms were evident when physiological symptoms first became significant (Fig. 2A). Inoculation of the more tolerant plants, M82, with mvF strain did not cause any visible or physiological symptoms. Nevertheless, the highly virulent vF strain did affect the M82 plants physiologically, as in all other cases, with infection being first detected as a change in transpiration, and later, as an effect on plant net mass (Table 1). Yet, no visible symptoms were detected over the course of the experiment.

At the end of every experiment, we documented the physical condition of the plants (Fig. 3A,B) and validated *F. oxysporum* f. sp. *lycopersici* infection through post-harvest analyses, including vascular browning assessment (Fig. 3C), fungal outgrowth tests (Fig. 3D), and PCR verification (Fig. 3E).The aforementioned assays provided evidence that the inoculation was successful and that the fungi colonized the inoculated plants independently of changes in physiological parameters.

**Fig. 3.**
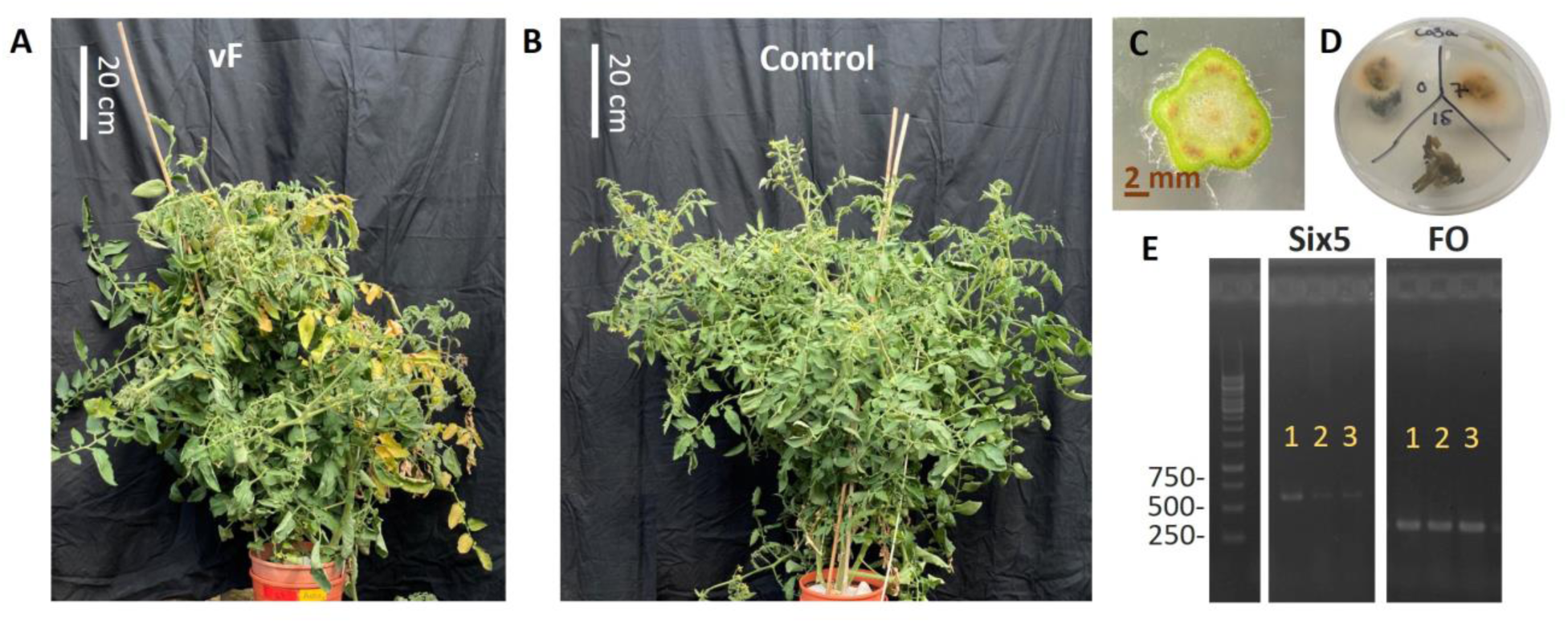
Tests to confirm infection. A, Visual symptoms of a *F. oxysporum* f. sp. *lycopersici* - inoculated Rehovot-13 plant, including wilting and typical chlorosis at 68 dai. B, Control Rehovot-13 plant. C, Browning of the vascular system in an inoculated plant, indicating fungal colonization. D, Fungal outgrowth from stem cutting of an inoculated plant observed on a PDA plate, slices are from different heights of the stem. E, Representative fungal outgrowth was subjected to PCR analysis to confirm the presence of *F. oxysporum* f. sp. *lycopersici* using the Six5 gene (667 bp) and FO ITS region (340 bp). In the figure are three samples from different plants marked in numbers, all samples have both six5 and FO genes.

### Quantitative ability of the PlantArray system: testing the system on fungal virulence and plant susceptibility levels

The selection of the *F. oxysporum* f. sp. *lycopersici* mvF and vF strains was validated through a standard seedling infection assay (Fig. S5). The seedlings infected with vF exhibited significantly more severe symptoms than those infected with mvF (*P* ≤ 0.03 ; Fig. S5). It is important to note that these assessments were terminal, as they required the physical cutting of plants for internal examination, and the severity of symptoms was subjectively scored in arbitrary units.

To evaluate the PlantArray system’s quantitative ability to assess the virulence of fungal strains, we used it to measure the effects of mvF and vF inoculation on susceptible R13 plants (Fig. 2A, B) . Twenty-six-days-old tomato plants were inoculated with either vF or mvF and grown under well-irrigated greenhouse conditions. Both *F. oxysporum* f. sp. *lycopersici* strains significantly reduced daily transpiration rates in the plants (*P* < 0.05, Fig. 2C), leading to wilt symptoms in the R13 plants (Fig. 2D,E). The impact of vF was more pronounced than that of mvF, with a noticeable reduction in transpiration seen as early as 16 dai, which was 6 days before similar observations in mvF-inoculated plants. This was coupled with a higher overall disease severity in the case of vF-inoculated plants (Fig. 2).

To assess the system’s ability to quantitatively evaluate tolerance levels of tomato cultivars to *F. oxysporum* f. sp. *lycopersici* (vF), we compared inoculated and non-inoculated R13 and M82 plants. The analysis of these quantitative parameters allowed us to calculate relative losses (as depicted in Fig. 4A,B). We found that M82 plants exhibited a relatively high level of physiological tolerance, experiencing only a 15% decrease in mass, as compared to non-inoculated controls. In contrast, the R13 plants displayed a more susceptible physiological response, with markedly pronounced losses, resulting in a 67% decrease in mass compared to controls. Additionally, we observed disparities in basic vigor traits between the two varieties, with the R13 plants exhibiting faster growth and accumulating more biomass than the M82 plants (Fig. 4B).

**Fig. 4.**
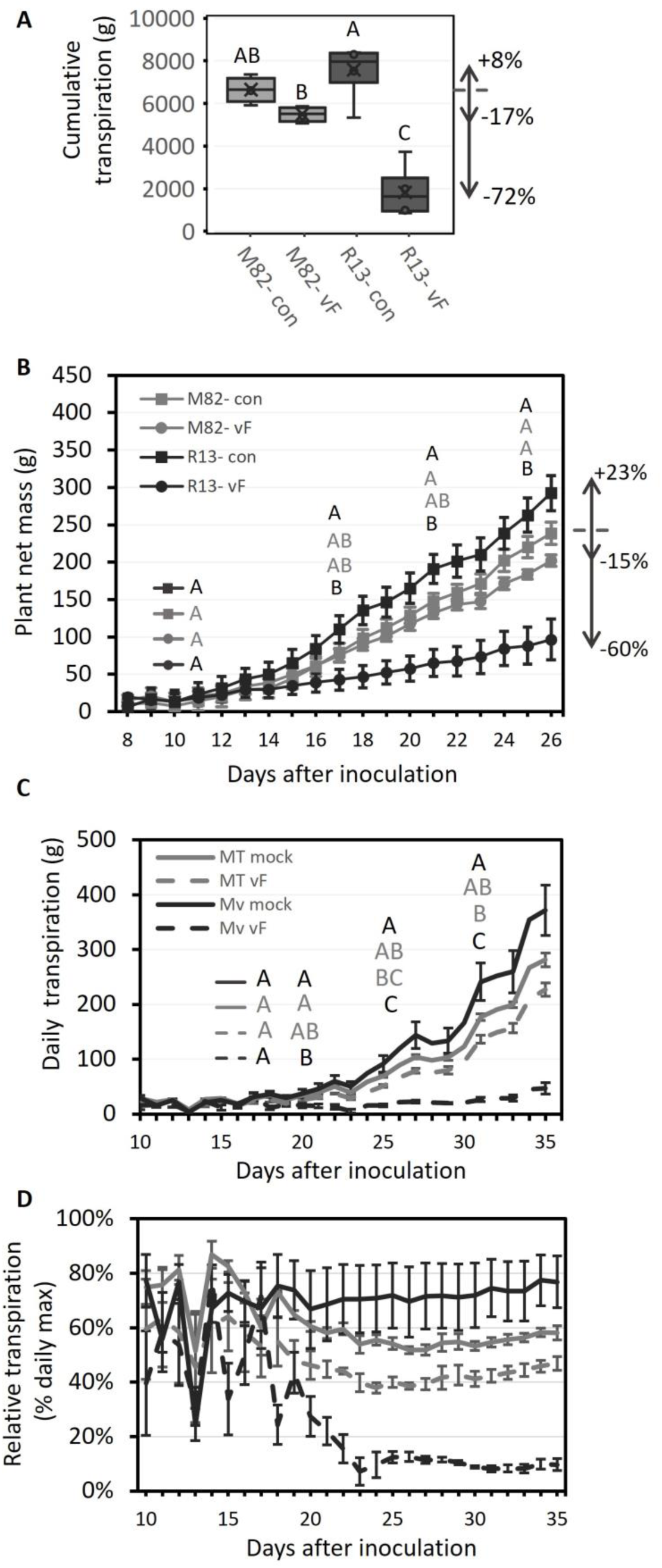
Differential impact of inoculation on water balance in resistant and susceptible plants: Profiling plant responses to *F. oxysporum* f. sp. *lycopersici* inoculation. Plants were inoculated with virulent *F. oxysporum* f. sp. *lycopersici* strains (vF) or mock-inoculated (control). A, This box-and-whisker plot represents the cumulative transpiration, which indicates the total water lost by the plants over the course of the experiment. Tolerant M82 plants are represented in gray, while susceptible Rehovot-13 (R13) plants are depicted in black. Arrows and percentages illustrate the relative differences compared to M82-control plants. B, Plant mass (mean ± SE) over the experimental period. M82 is shown in gray; R13 is shown in black; circles indicate inoculated plants, while squares indicate non-inoculated, control plants. Letters indicate statistically significant differences between groups (*P<*0.05), and their placement corresponds to the first instance of a significant change during the experiment. The differences between each set of plants and the M82-control (dashed line) at the end of the experiment are presented as arrows and percentages. C, Daily transpiration (mean ± SE) over the experimental period. Resistant cv. Motelle (MT) plants are shown in gray and susceptible cv. Marmande Verte (Mv) plants are shown in black. Dashed lines indicate inoculated groups. Using the same data, we calculated the (d) daily transpiration relative to the daily maximum values (mean ± SE). Different letters indicate statistically significant differences (P = 0.05; Tukey-Kramer test).

Subsequent tests were carried out on various tomato cultivars, to evaluate the efficacy of the system across varieties differing in their levels of resistance to *F. oxysporum* f. sp. *lycopersici* . We specifically examined the performance of cv. Motelle (MT), a cultivar known to be highly resistant to *F. oxysporum* f. sp. *lycopersici*, cv. Moneymaker (MM), which is nearly isogenic to MT yet susceptible to *F. oxysporum* f. sp. *lycopersici*, and the *F. oxysporum* f. sp. *lycopersici* susceptible cultivar Marmande Verte (MV). Consistent with our earlier findings, we observed a reduction in transpiration in infected MV plants (Fig. 4C). Moreover, the decrease in transpiration, expressed as a percentage of the daily maximum transpiration (Fig. 4D), provided a clear and quantitative measure of the functional losses among the different groups. Specifically, the MV plants infected with vF transpired only 10% of the daily maximum.

*F. oxysporum* f. sp. *lycopersici*-inoculated MM plants transpired significantly less and also gained less biomass than non-inoculated MM plants (*P* < 0.05; about 20% less transpiration; Fig. S6). Non-inoculated MM plants showed a trend of increased productivity relative to MT plants, but that difference was not significant (Fig. S6). As expected, the resistant MT plant did not exhibit a significant change in mass or transpiration despite *F. oxysporum* f. sp. *lycopersici* inoculation.

### Impact of *F.oxysporum* f. sp. *lycopersici* filtrate on leaf hydraulic conductance (K_leaf_)

Reduction in transpiration due to *F. oxysporum* f. sp. *lycopersici* infection may be due to fungal hyphae clogging the xylem vessels, and/or to a signal transduction pathway that reduces hydraulic conductance at an early stage of infection. To assess the effect of isolated culture-filtrate toxins on leaf hydraulic conductance (K_leaf_), we compared the impact of fungal culture filtrate, perfused into the leaf vascular system. As previous studies revealed that chitin and abscisic acid (ABA) both decrease leaf hydraulic conductance (K_leaf_) by reducing the osmotic water permeability (P_f_) of bundle-sheath cells, we used these compounds as control treatments in our experiments. Both the susceptible MM and the resistant MT plants exhibited similar reductions in K_leaf_ when exposed to control stress-treatments of chitin and ABA (Fig. 5A; 48% and 89%, respectively). These results are consistent with findings reported in previous studies (Attia et al. 2020). However, the susceptible MM cultivar showed a significantly greater decrease in K_leaf_ in response to the highly toxic-treated group (vF; 65%), as compared with the moderately toxic group (mvF; 42%), and a significant reduction in K_leaf_ as compared to the MT cultivar in response to both of these treatments (Fig. 5B). Specifically, MT plants showed a 25% reduction in K_leaf_ when infected with mvF and a 35% reduction when infected with vF. This decrease in K_leaf_ suggests a strong response to the presence of fungal toxins, which may implicate these toxins in the mechanism that reduces K_leaf_.

**Fig. 5.**
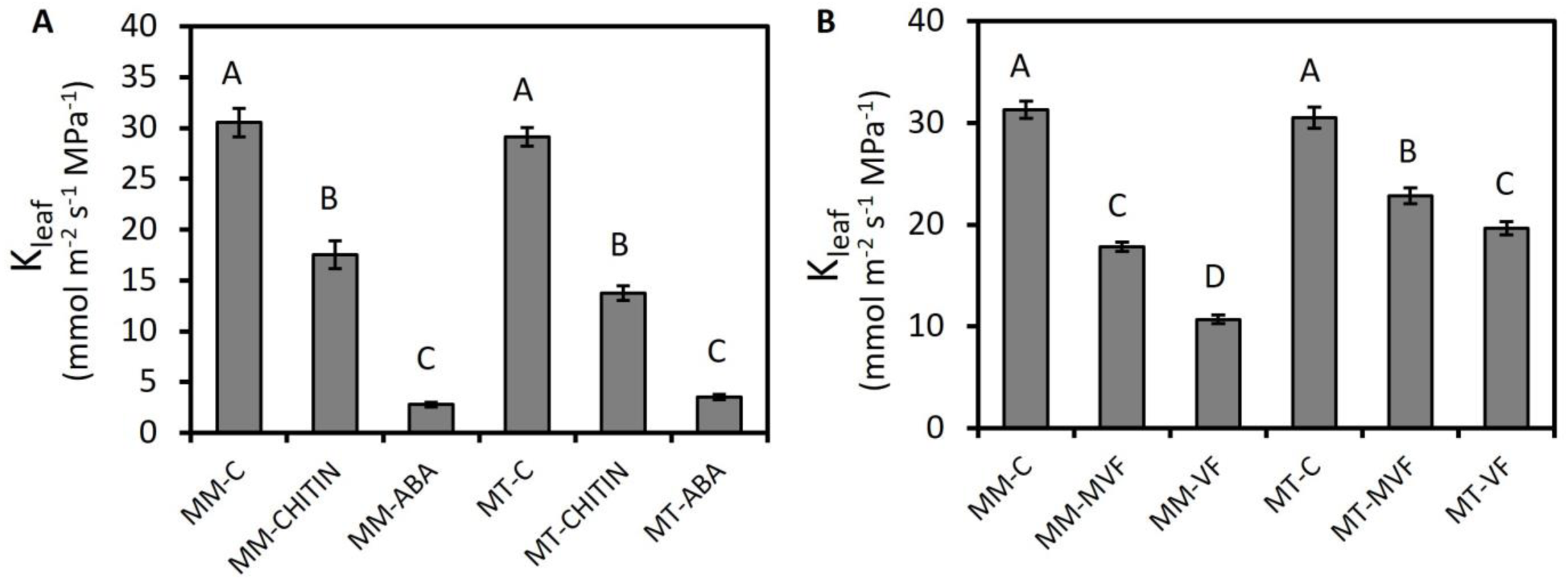
Effects of chitin, ABA, and fungal toxins on leaf hydraulic conductance (K_leaf_). Detached leaves from cv. Moneymaker (MM) and cv. Motelle (MT) tomato plants were treated for 2 to 4 h with (A) chitin (0.2 mg/mL) or ABA (10 μM), or (B) toxins released to the medium during fungal growth from both moderately virulent and virulent *F. oxysporum* f. sp. *lycopersici* strains. Control groups were treated with 0.5% of the control filtrate. MVF and VF: Treated with 0.5% of the culture filtrate from the moderately virulent and virulent *F. oxysporum* f. sp. *lycopersici* strains, respectively. Different letters indicate significant differences between treatments according to the Tukey-Kramer HSD test (*P* < 0.05). Data points are means (± SE) from three to five distinct experiments, including 12 to 30 biological repetitions.

Additionally, we observed substantial reductions in the leaf transpiration rate (E) and leaf water potential (Ψleaf) following treatment with stress-controls chitin and ABA in both MT and MM plants (Fig. S7). Furthermore, similar to our whole-plant results, the MM cultivar exhibited a significant reduction in leaf E under the mvF treatment, which became even more pronounced under the vF treatment (24% and 43%, respectively). In contrast, the MT cultivar did not show any reduction in leaf E under the mvF treatment and its reduction of E under the vF treatment was less pronounced than that of the MM plants (18%, Fig. S8).

### Prediction of disease symptoms in infected plants under dynamic environment

To identify the factors influencing the occurrence of disease symptoms in infected plants, we examined various internal and external factors (Table S1). Our automated system efficiently collected this information, enabling the generation of a detailed dataset encompassing environmental conditions, infection parameters, and plant properties. We entered these data into a model to predict the occurrence of disease events with symptoms. Specifically, we focused on the interaction between cultivar and fungal strain, as well as plant properties on the date of inoculation, such as mass and age. Additionally, we examined the impact of the environmental parameters at 3 dai that were automatically collected from the PlantArray system, including temperature, relative humidity (RH) and daily light integral (DLI). We only included experiments in our analysis where the infected plants showed fungal outgrowth at the post experiment analysis, indicating successful infection (Table S1, “Fungi test”). By employing logistic regression analysis and a backward elimination approach, we identified the best model for predicting whether disease symptoms would appear (Model S1). The model demonstrated a high degree of accuracy (*R*^2^ = 1, *P* < 0.0001). The significant variables in this model include the interaction between the plant cultivar and fungal strain (*P* = 0.011), plant initial mass (*P* < 0.001), RH (*P* < 0.001), and DLI (*P* < 0.001). Furthermore, our findings suggest that smaller plant mass (coefficient of -52.64, not significant), higher RH (coef. 58.24, n.s.), and higher DLI (coef. 34.1, n.s.) during the initial 3 days are associated with an increased likelihood of the development of disease symptoms in infected plants. However, it is important to note that the observed coefficients had high chi-square values (ChiSq = 0), but lacked statistical significance (*P* = 0.99), likely due to the limited number of repetitions within each plant–pathogen group.

Testing the model on a newly produced experiment with the R13 tomato cultivar and both mvF and vF fungal strains, with small plant mass (3.6 g), during the first 3 days, the average RH was 61.19% and the DLI was 19 mol/(m²*day). The model predicted the occurrence of symptoms (Probability of "yes" symptoms: 1), which indeed appeared by the second week of the experiment.

### Testing the method generalizability in the potato*–Phytophthora infestans* pathosystem

To evaluate the applicability of the PlantArray system in detecting early disease symptoms in non-vascular plant-pathogen interactions, we tested its potential in potato plants inoculated with *P. infestans*. The initial symptoms of this disease are irregular spots that are light to dark green in color, moist in appearance, and necrotizing. These lesions appeared 4 to 5 dai (Fig. 6E, day 7). The daily transpiration and plant net mass were significantly different at 5 dai (Fig. 6A,B). However, it took only 3 days from inoculation for significant differences in transpiration and E (transpiration normalized to mass) at midday to develop between the control and the infected plants (*P* < 0.05, Fig. 6C,D).

**Fig. 6.**
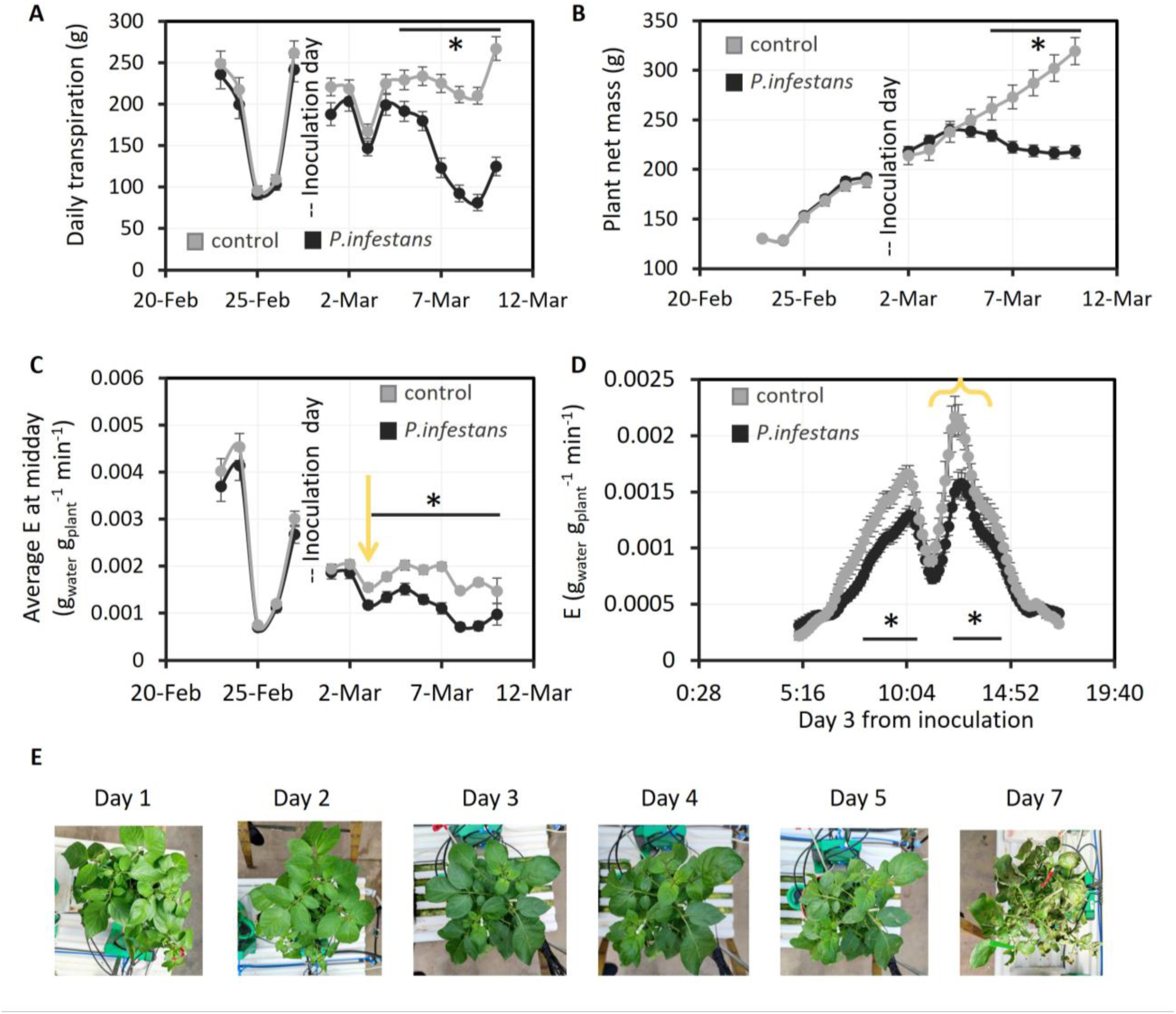
Quantitative early detection of *Phytophthora infestans* in potato. Potato plants inoculated with *P. infestans* were compared with control (mock-infected) plants. Inoculation day was 28-Feb; Day 1 after inoculation was 1-Mar. A, Daily transpiration of the whole plants, means ± SE. B, Plant net mass over the entire experimental period, means ± SE. C, Daily average E between 11:00 to 14:00. At this time of day, E is usually at its highest. D, E values throughout the third day from inoculation, means ± SE. The orange arrow and bracket are pointing at the same data represented as point or continues data, respectively. Each treatment had 18 plants, asterisks indicate significant differences between the inoculated and control groups (*t*-test; *P* < 0.05). E, Pictures of inoculated plants at different points in time after inoculation.

## Discussion

Visual estimation is the main method used to detect and assess plant diseases. However, the fact that visual estimation often relies on subjective assessment and the late appearance of visible symptoms limit the efficiency of that approach. To overcome these limitations, we present a physiological functional method that provides objective and quantitative measurements of disease progression, enabling both early detection and precise measurements for research applications. This method allowed us to (i) detect *F. oxysporum* f. sp. *lycopersici* infection at an early stage, before the appearance of visual symptoms; (ii) quantify the virulence and pathogenicity levels of two *F. oxysporum* f. sp. *lycopersici* strains, as well as (iii) to compare the susceptibility levels of different tomato cultivars.

### Early detection of disease

Our findings suggest that transpiration serves as an early indicator of *F. oxysporum* f. sp. *lycopersici* and *P. infestans* infection (Fig. 6). Notably, in the case of *F. oxysporum* f. sp. *lycopersici*, this indication came up to 49 days before the manifestation of visual symptoms (Table 1, Row 5) and an average of 35.2 days before any morphological symptoms were observable. Other studies have reported a decline in vascular flow or water loss slightly before the appearance of wilt symptoms in infected plants (Feng et al. 2022; Street and Cooper 1984; Wang et al. 2012). To document this, Wang et al. (2012) and Feng et al. (2022) used a Li-6400 gas exchange system (Li-Cor Inc.); whereas Street and Cooper (1984) used a Scholander pressure bomb. In contrast to the simple assessment using the PlantArray system, the aforementioned methods are time-consuming and laborious.

### Quantification of disease severity: host susceptibility and pathogen virulence

Our study provides a new approach for quantifying *F. oxysporum* f. sp. *lycopersici* aggressiveness in tomato plants by using transpiration decrease as a comparative test for *F. oxysporum* f. sp. *lycopersici* virulence. We found that vF caused a significantly greater decrease in transpiration than mvF (Fig. 2C), indicating a higher level of pathogen aggressiveness. This was confirmed by traditional methods such as visual assessment of plants when symptoms appeared (Fig. 2D) and visual ratings of the level of infection in seedlings (Fig. S5). However, visual ratings of *F. oxysporum* f. sp. *lycopersici* were challenging due to the difficulty in comparing, assessing, and scoring the disease severity. Stewart and McDonald (2014) demonstrated the divergences between scorers’ visual assessments and the accurate values (Stewart andand McDonald 2014). Other researchers have also recognized the need for quantitative measures of pathogen aggressiveness, as visual estimation alone is insufficient (Bock et al. 2020; Gale et al. 2003; Ilgen et al. 2009). This study introduces an approach utilizing whole-plant transpiration and continuous biomass assessments for assessing pathogen virulence, providing a potentially valuable tool for comparative testing of *F. oxysporum* f. sp. *lycopersici* strains.

The literature defines the M82 tomato as moderately resistant (Sela-Buurlage et al. 2001) or tolerant to *F. oxysporum* f. sp. *lycopersici* race 2; whereas R13 is known to be susceptible to *F. oxysporum* f. sp. *lycopersici* race 2 (Sarfatti et al. 1989). Despite these dichotomic definitions used to categorize a plant as either tolerant or susceptible to a pathogen, there remains a need for more quantitative, reliable and effective measurement techniques to improve our understanding of tolerance levels (as reviewed by Robb 2007). This review suggest that quantitative methods would better capture tolerance levels, aiding breeding and plant-pathogen research. Additionally, for vascular wilt disorders, disease severity is usually scored in a semi-quantitative fashion using a disease index (Robb 2007), as presented in Figure 1E. Although measuring disease severity can be challenging, yield is considered less controversial and is a reliable test for plant health (Scott 2005). Our functional phenotyping method allows the classification of resistance to susceptibility on a percentage scale, as opposed to a dichotomous terminology. For instance, infected M82 plants exhibited a 15% decrease in plant mass; whereas infected R13 plants exhibited a mass decrease of 67% (Fig. 4B). Thus, the quantitative, objective nature of our functional phenotyping method significantly refines the evaluation of plant tolerance level, which could provide valuable insights into cultivar–pathogen dynamics and the risks and opportunities associated with those dynamics.

We posit that our findings likely correlate with plant performance in agricultural settings. Transpiration is intrinsically linked with CO_2_ absorption, photosynthesis, plant growth and yield. The correlation between transpiration and crop yield is complex, as both factors are influenced by a multitude of environmental and physiological processes. Nevertheless, in many modern crops, whole-plant transpiration has been found to be linearly correlated with yield (Gosa et al. 2019). This relationship is particularly evident in tomato (Chaka Gosa et al. 2022), indicating that our functional phenotyping method may serve as a reliable prediction for productivity in that crop.

### The cost of resistance

Constitutive activation of defense mechanisms might negatively affect plant productivity in the absence of pathogen infection (Zhao et al. 2017). In our experiments, non-inoculated controls showed a resistance cost: M82 had 19% less biomass than R13, and MT had 24% less transpiration than Mv (Fig. 4B, C). A similar resistance penalty was reported by Shteinberg et al. (2021), who found that tomato lines that were susceptible to *Tomato yellow leaf curl virus* performed better under optimal conditions than a resistant line (Shteinberg et al. 2021). Defense mechanisms divert energy from growth, making less-defended plants more productive in the absence of pathogens (Cipollini et al. 2014; Herms and Mattson 1992). We suggest that this quantification may be useful for breeders and farmers, helping them to fine-tune their selections and to invert their dichotomic scope of resistance-susceptible terminology to levels of productivity, risk, and cost.

### Impact of fungal toxins on plant hydraulics

The decrease in the transpiration of the infected plants could be due to hydraulic blockage resulting from physical clogging of the xylem by insoluble fungal materials, such as spores, mycelia, and polysaccharides. It also could be due to a signaling response pathway activated by a soluble fungal toxin that may have traveled through the vascular system to the leaves. Our observation that transpiration, K_leaf_, and water potential were all reduced following the exposure of leaves to fungal filtrate (Fig. 5) highlights the considerable impact of the signaling pathway on the plant’s overall water balance. Researchers have tested the effects of the crude fungal toxins on wilting (Madhosingh 1995), the death of leaf protoplasts (Sutherlandt and Pegg 1992) and callus growth in culture media (Scala et al. 1985). This study examined the effect of fungal toxins on leaf hydraulic conductance, offering insights into the impact of signaling pathways on the plant’s overall water balance. Our findings suggest that the signaling pathway plays a crucial role in the plant’s early response, affecting it even before any physical clogging occurs.

Interestingly, the vF toxins secreted into the medium had a more severe effect than the mvF toxins and leaves with high levels of immunity were less affected than susceptible leaves. This aligns with previous research involving culture filtrates, in which plant lines capable of tolerating or resisting toxins as seedlings or protoplasts also exhibited resistance under greenhouse conditions (Hartman et al. 1984; Mcleod and Smith 2012; Sutherlandt and Pegg 1992). Therefore, the impact of these toxins on plant hydraulics is similar to the effect of fungal infection.

### Disease symptoms in infected plants under dynamic environment

Our research, conducted in greenhouse conditions, underscores the significant effects of environmental factors and plant–fungus interactions in disease progression in the *F. oxysporum* f. sp. *lycopersici* –tomato pathosystem. Through analyzing multiple experiments (Table S1), we found that smaller plants are more likely to exhibit wilting symptoms within 30 to 70 days post-inoculation, and environmental factors like high DLI and high RH promote disease development (Model S1). These findings highlight the importance of considering early post-infection environmental factors and plant–fungus interactions when predicting disease progression.

In 11 out of 19 experiments we did not observe any symptoms, neither visual nor physiological. The lack of disease symptoms in inoculated plants is not uncommon. Some studies report late symptoms exceeding 60 days, particularly in experiments conducted under field-like conditions (Paugh and Swett 2020; Rekah et al. 2001). Plants under sub-optimal environmental conditions are more susceptible to pathogen infections due to weakened defense mechanisms, while optimal conditions generally enhance plant resistance but may also facilitate pathogen spread under certain circumstances (Atkinson and Urwin 2012). Optimal conditions for greenhouse tomato cultivation include daytime temperatures of 21 to 27°C and nighttime temperatures of 18 to 20°C, with relative humidity levels of 60 to 85% (DryGair n.d.; Shamshiri et al. 2018). Tomato growth requires a minimum photosynthetic photon flux density (PPFD) of 200 to 300 μmol/m² /s¹, with an optimal level of 600 μmol/m²/s¹ or higher (Klaring and Korner 2020). Furthermore, maintaining a vapor pressure deficit (VPD) between 0.3 to 1.0 kPa (Grange and Hand 1987; Shamshiri et al. 2018), ensuring adequate ventilation, and carefully managing fertigation are critical for healthy plant growth and development. In fact, in many experimental systems the conditions are set to see the strongest symptoms, often because it is very hard to distinguish more subtle symptoms. Our growth facility, optimized for tomato cultivation, may have contributed to enhanced defense mechanisms in the plants, potentially enabling them to better withstand pathogen challenges. In this set-up, then, measuring quantitatively plant disease brings our investigation closer to the agricultural setting.

### Limitations and future research

The physiological functional phenotyping method described here faces three key limitations: (i) its reliance on pot-based experiments in greenhouse settings, (ii) the perceived high costs of phenotyping technologies, and (iii) the restricted applicability of high-throughput methods in field conditions. However, pot-based phenotyping provides controlled, dynamic, and accurate simulations of plant responses, allowing for precise environmental manipulation and rapid data collection, which can ultimately save time and resources. Moreover, pot-based system like the one we present here is expected to reveal markers for early infection that could in the future be translated to field-based discovery platforms. Future research should couple physiological measurements with spectrum-based analysis. Regarding costs, the investment is relatively manageable; for example, the budget for a single gas exchange system could support around 30 lysimeter systems. These lysimeters continuously measure whole-plant gas exchange and provide irrigation feedback, significantly improving the efficiency of phenotypic assessments. Additionally, technological advancements have led to the development of large-scale lysimeter systems that are now functional in field settings. While field lysimeters have higher upfront costs, their integration with field sensors offers substantial potential for bridging the gap between controlled greenhouse experiments and real-world agricultural applications.

Given that both plant and microbial development are influenced by dynamic environmental conditions, diagnostic approaches that rely on arbitrary time points post-inoculation (e.g., molecular techniques such as RNA sequencing (RNA-seq) and sap flow analyses) may not accurately reflect the physiological state of the plant during infection and have shown significant differences within several hours (Guo et al. 2021). Thus, it is crucial to carefully choose the time points for sample collection, emphasizing our physiological benchmark approach for determining the optimal sampling times. By standardizing sample collection based on specific physiological parameters, such as an 80% reduction in transpiration compared to control plants, we can ensure that molecular analyses are conducted when the host plant is in a defined state of disease progression. This approach is expected to improve the accuracy of molecular data and also enhance the reproducibility of experiments across different studies. As demonstrated in the study by Fox et al. (2018) on *Pinus*, the simultaneous continuous monitoring of whole-plant physiological parameters with molecular studies enables a comprehensive understanding of both. Our approach opens the way to conduct molecular analyses before visual symptoms are observed but right after an effect of the pathogen on plant physiology is observed thus pinpointing early molecular events in pathogenesis.

While this study focused on the vascular pathogen *F. oxysporum* f. sp. *lycopersici* and to a lesser extent on the foliar pathogen *P. infestans*, further research should explore the applicability of this system to other pathosystems, including bacterial, viral, and nematode pathogens. Our results demonstrate that pathogens affecting the vascular system can cause a reduction in transpiration rate, and we also observed that transpiration can decrease due to mycotoxin exposure. Fungal and bacterial toxins are common effectors of plant diseases and therefore the system presented here may be useful to study their effect. Additionally, testing the interactions between abiotic and biotic stresses, as well as co-infections, could provide a broader understanding of the system’s potential for screening diverse diseases across crops.

## Conclusions

Our study revealed that a high-throughput physiological monitoring system is suitable for the early detection and quantification of Fusarium wilt disease in tomato caused by *F. oxysporum* f. sp. *lycopersici*. Fundamentally, we found that whole-plant transpiration is a reliable indicator of plant health. Furthermore, the examined system also allows for simultaneous real-time analysis, which enables a quality comparison between pathogen virulence and host susceptibility. Indeed, the high-throughput monitoring of the physiological responses of tomato plants following infection with *F. oxysporum* f. sp. *lycopersici* revealed a spectrum of plant–pathogen interactions. This system may help breeders and researchers to evaluate plant tolerance and pathogen virulence on a quantitative scale, thereby making plant– pathogen research more efficient and more accurate. Further studies should be done to assess whether this system may be applied to other pathosystems. In addition, this approach generates large, annotated datasets of plant–pathogen–environment interactions, which could potentially be integrated into computer-based models for predicting plant reactions and estimating plant losses using machine learning and artificial-intelligence tools. Overall, this non-destructive, high-throughput method for monitoring plant health and disease progression has great potential for improving research in the field of phytopathology. Additionally, our study demonstrates the applicability of this physiological monitoring system for early detection and quantification of *P. infestans* infection in potato, offering a versatile tool for plant-health assessment in diverse pathosystems.

## Acknowledgments

This research was supported by the Shoenberg Research Center for Agricultural Science (Grant #3175006230) and the Israel Science Foundation (Grant No. 1043/20). We extend our gratitude to Yael Rekah from the Hebrew University, Israel, for kindly providing ‘Marmande Verte’ and ‘Motelle’ seeds originally sourced from H. Laterrot, INRA, France. Special thanks to Dr. Yaniv Rotem (Hazera Genetics) for supplying *F. oxysporum* f. sp. *lycopersici* strain fr2T and offering valuable advice. We also acknowledge Nofit Vaknin,Phytopathology Coordinator (Hazera seeds), for her insights into traditional screening methods.

## Supporting Information

**Fig. S1** Weather properties continually monitored in the greenhouse.

**Fig. S2** Experiment design

**Fig. S3** Visual Disease Severity Scoring Scale

**Fig. S4** A classic seedling disease assay confirmed the resistance of MT and the susceptibility of MM, R13 and M82 to *F. oxysporum* f. sp. *lycopersici* .

**Fig. S5** A classic seedling disease assay confirmed that vF is more virulent than mvF.

**Fig. S6** Plants of susceptible (MM, gray) and resistant (MT, black) near-isogenic lines were inoculated with vF (dashed line).

**Fig. S7** Effects of chitin and ABA on the hydraulics of leaves detached from MM and MT plants.

**Fig. S8** Effects of toxins released from the moderately virulent and virulent *F. oxysporum* f. sp. *lycopersici* strains on leaves detached from MM and MT tomato plants.

**Table S1** Summary of experiments with different *F. oxysporum* f. sp. *lycopersici* strains and plants of varying levels of susceptibility.

**Model S1** Logistic regression model of symptom occurrence.

