## Supporting Information for "Early Detection and Quantification of Fusarium Wilt in Greenhouse-Grown Tomato Plants Using Water-Relation Measurements"

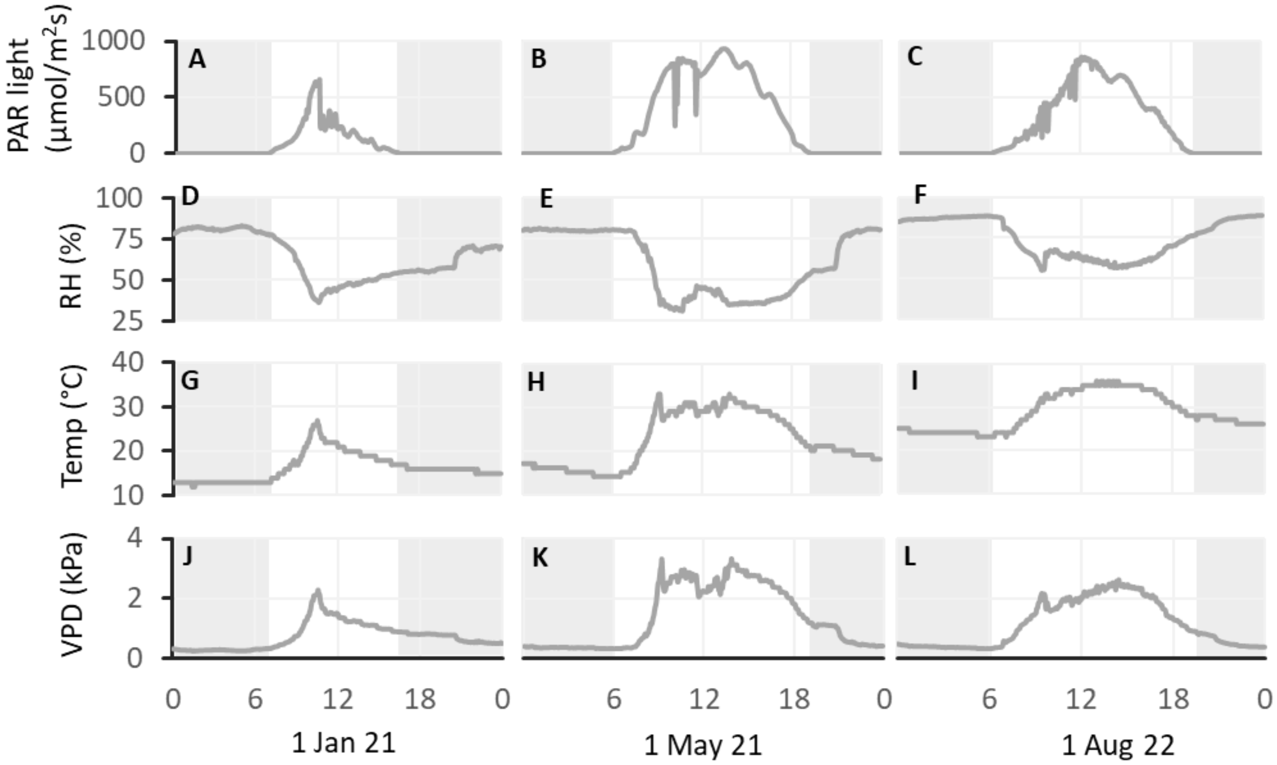


**Fig. S1 Weather properties continually monitored in the greenhouse.**

Representative daily measurements at three dates: 1 January 2021, 1 May 2021 and 1 August 2022. Daily continuous values of (A to C) PAR light, (D to F) relative humidity (RH), (G to I) temperature and (J to L) vapor pressure deficit (VPD). Nighttime hours are shaded.

**Fig. S2 Experiment setup and design**

In our greenhouse pots were arranged on tables, with each table holding 12 pots. Treatments were allocated using a randomized block design within each table to reduce spatial variability and ensure an unbiased distribution of treatments across the greenhouse. Each treatment was replicated 4 to 6 times throughout the tables to ensure statistical validity. The randomized design shown here corresponds to Experiment 2, as detailed in Table 1, at that experiment only 30 out of the 60 PlantArray units were used.

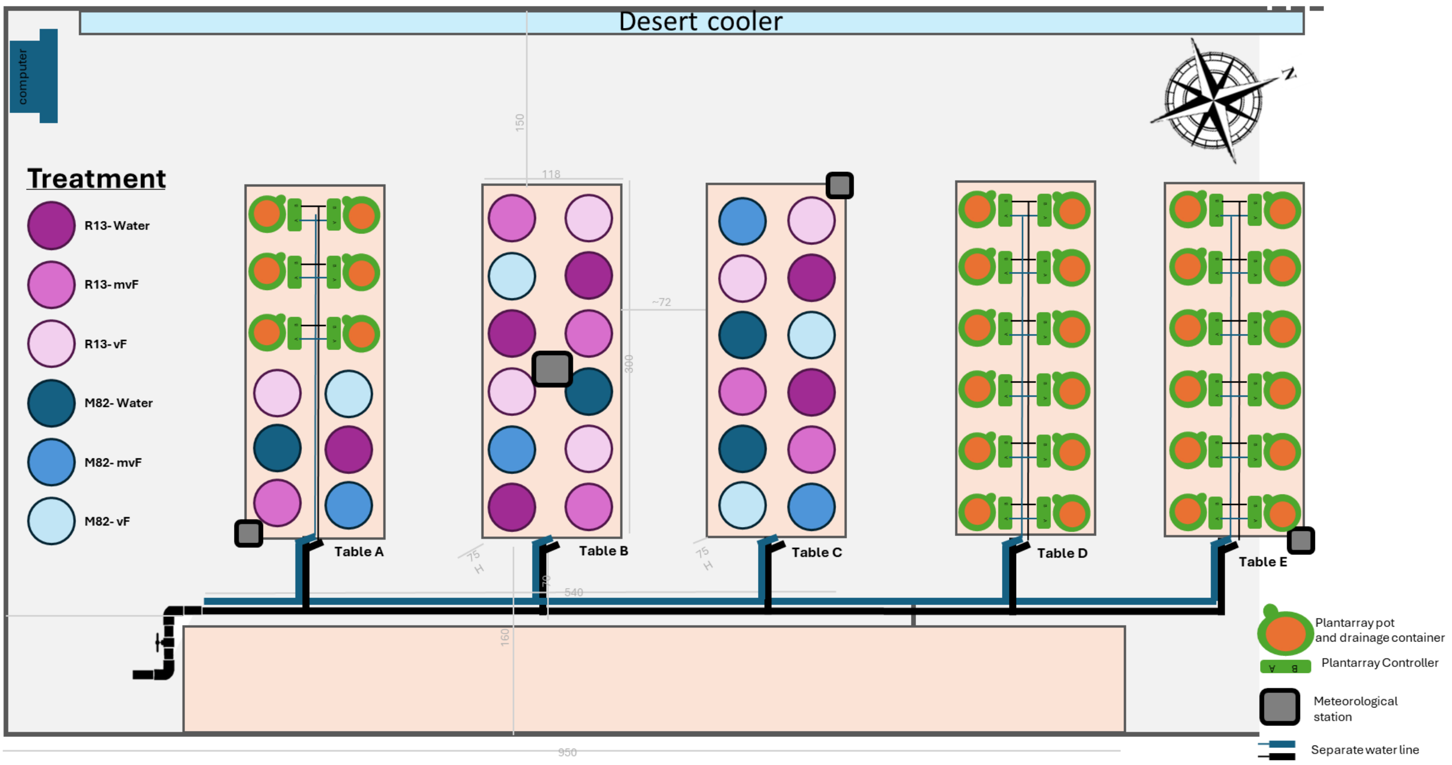

**Fig. S3 Visual Disease Severity Scoring Scale**

Photographic illustrations of the visual disease severity scoring scale used to monitor Fusarium wilt progression in tomato plants. The scoring scale ranges from 0 to 4, with 0 representing asymptomatic plants with no visible symptoms, 1 indicating weakly symptomatic plants with less than 25% of leaves showing chlorosis or wilting, and 2 representing moderately symptomatic plants with 25–50% of leaves affected. A score of 3 denotes highly symptomatic plants with more than 50% of leaves wilted, though the plants remain alive, while a score of 4 represents dead plants. This figure provides a visual reference for the disease severity levels described in the study.

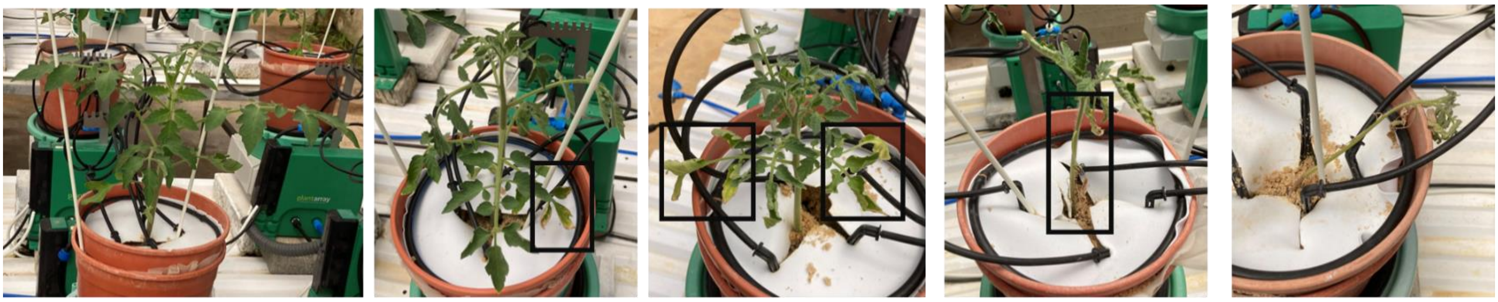

0 = asymptomatic

1 = weakly symptomatic (<25% of leaves chlorotic or wilted)

2 = moderately symptomatic (25–50% of leaves were chlorotic or wilted)

3 = highly symptomatic (>50% of the leaves wilted, but plants were alive)

4 = dead

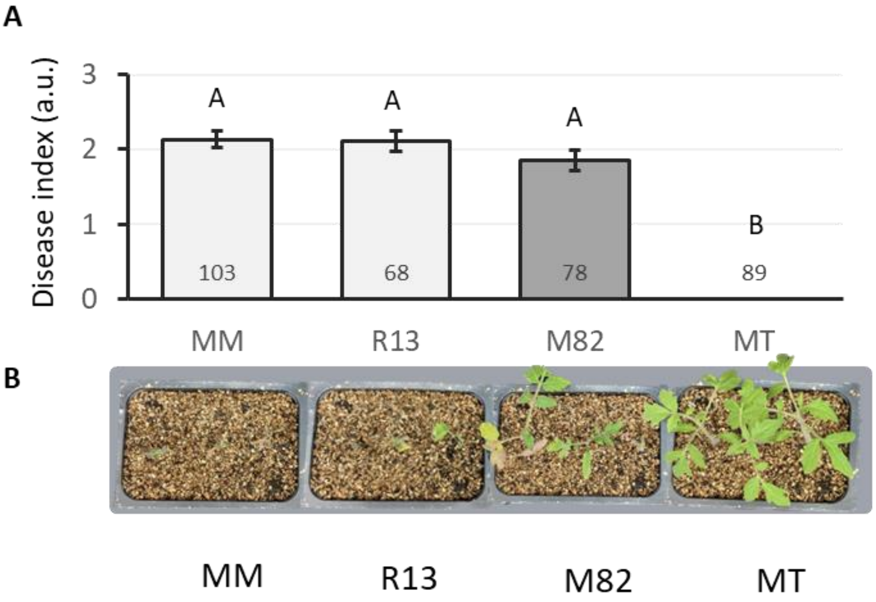


**Fig. S4 A classic seedling disease assay confirmed the resistance of MT and the susceptibility of MM, R13 and M82 to *F.oxysporum f. sp. lycopersici*.**

Severity of disease in tomato seedlings inoculated with *F.oxysporum f. sp. lycopersici*. MT (resistant), M82 (tolerant), MM (susceptible) and R13 (susceptible) plants were inoculated with *F.oxysporum f. sp. lycopersici*, race 2. A, Severity of disease in tomato seedlings was assessed at 21 days after inoculation using a disease index based on arbitrary units (a.u.). Two independent experiments were conducted, each including two independent *F.oxysporum f. sp. lycopersici* strains, with approximately 20 plants per batch. The total numbers of plants are presented in the bars. Different letters indicate statistical differences between groups, according to the Tukey-Kramer test. B, Representative photographs of plants infected with vF.

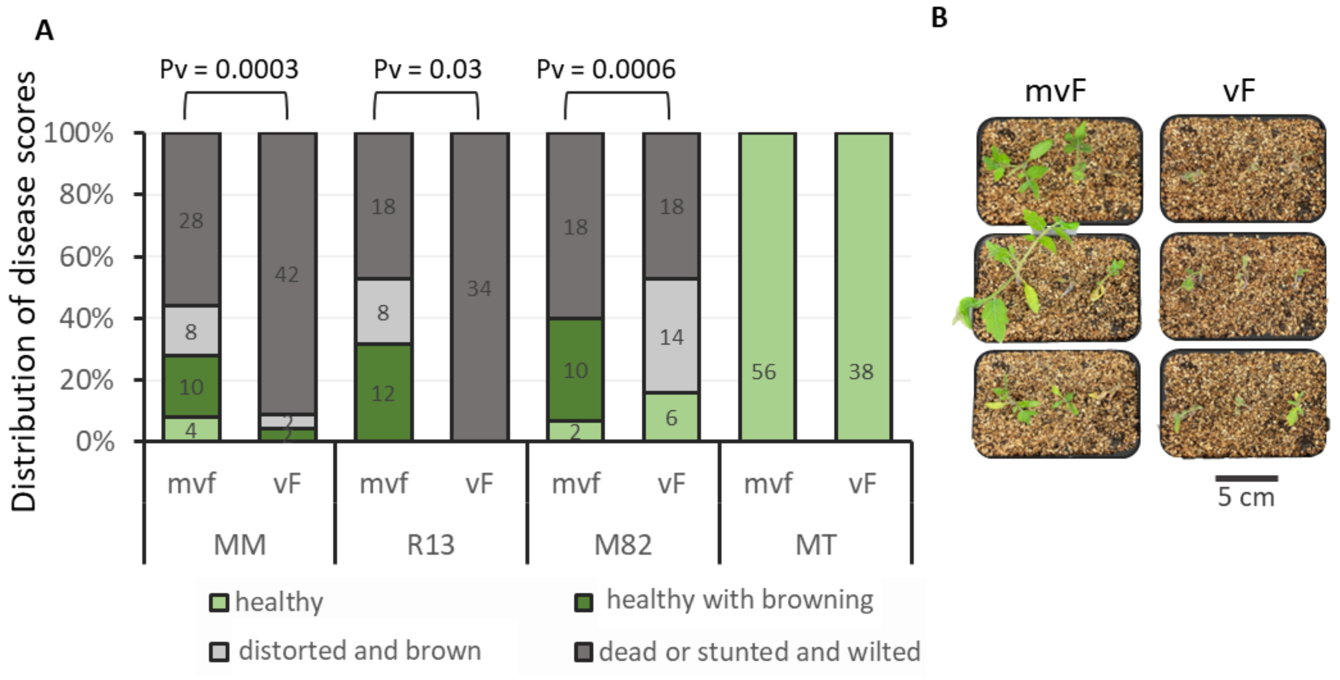


**Fig. S5 A classic seedling disease assay confirmed that vF is more virulent than mvF.**

Twelve-day-old seedlings of MT (resistant), M82 (tolerant), MM (susceptible) and R13 (susceptible) tomato plants were inoculated with *F.oxysporum f. sp. lycopersici*, race 2. Plants were inoculated with either a moderately virulent strain, mvF or the highly virulent strain, vF. Control plants were mock-inoculated. Disease severity was assessed 21 days after inoculation using a disease index based on arbitrary units (a.u.; see Materials and Methods). A, Disease distribution differed between the tomato variates and within the *F.oxysporum f. sp. lycopersici* treatments. Probability values were obtained using the chi-squared likelihood-ratio test, to identify significant differences between the levels of disease caused by the different *F.oxysporum f. sp. lycopersici* strains. B, Representative pictures of MM plants infected with mvF or vF, 3 plants in each pot.

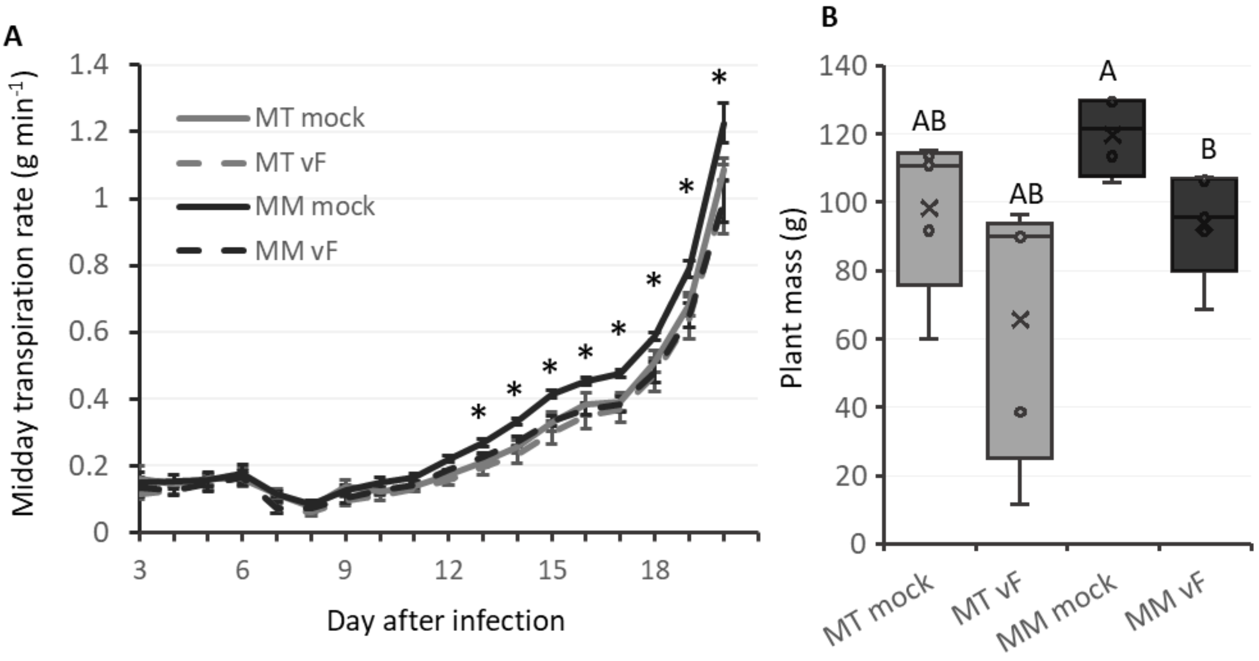


**Fig. S6 Plants of susceptible (MM, gray) and resistant (MT, black) near-isogenic lines were inoculated with vF (dashed line).**

A, Average transpiration rate at midday (between 10:00 to 14:00) ± SE. Asterisks denote significant differences between MM mock-inoculated and MM vF-inoculated plants (*t*-tests; $*P \leq 0.05$); 4–5 plants in each treatment. B, Box-and-whisker plot represents the plant mass at the end of the experiment (Day 20 after inoculation). Different letters indicate statistically significant differences (Student's *t*-test, $P = 0.05$).

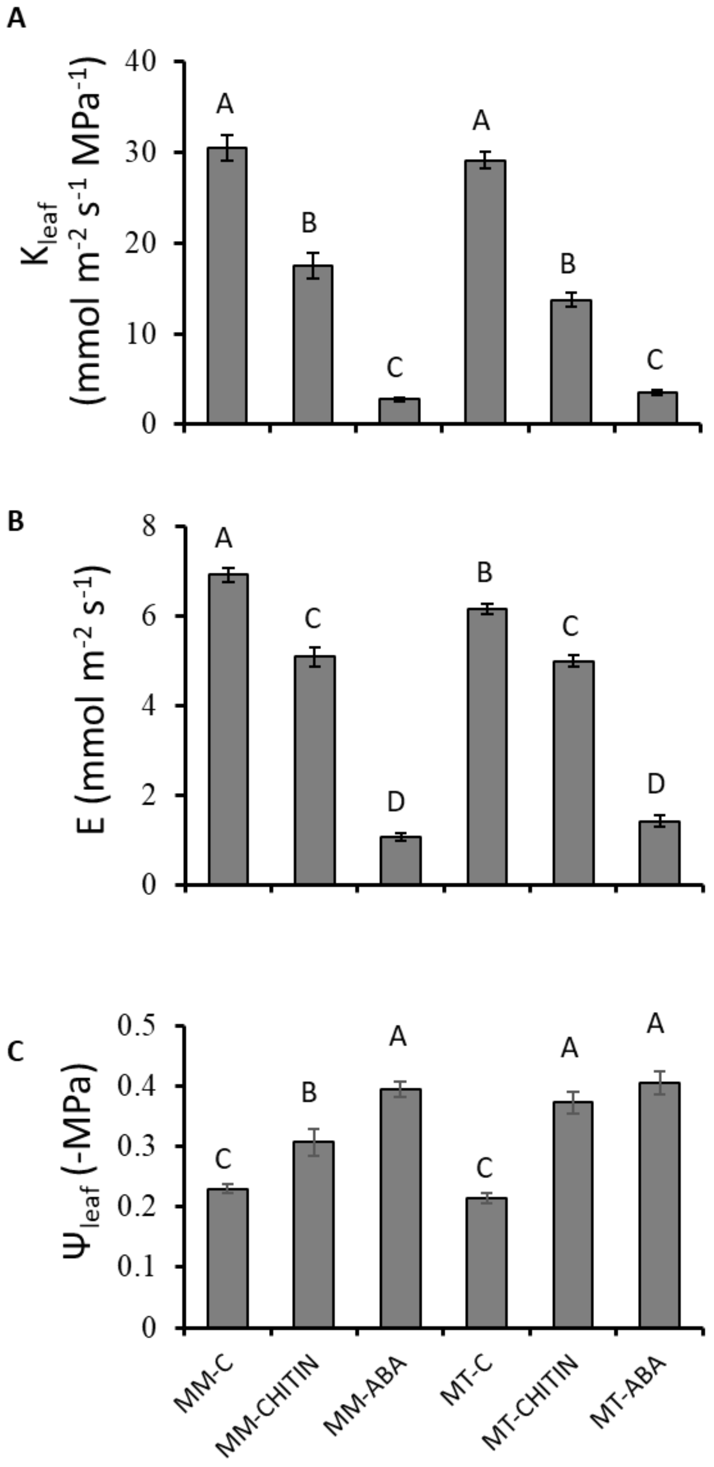


**Fig. S7 Effects of chitin and ABA on the hydraulics of leaves detached from MM and MT plants.**

A, Leaf hydraulic conductance, $K_{leaf}$, B, transpiration rate (E) and C, leaf water potential, $\Psi_{leaf}$ after 2–4 h of the xylem-fed chitin and ABA treatments. A, is identical to Fig. 5. C: Control plants fed with AXS alone. CHITIN: Treated with 0.2 mg/ml of chitin. ABA: Treated with 10 µM ABA. Different letters indicate significant differences between treatment results based on the Tukey-Kramer HSD test ($P < 0.05$). Data points are means (± SEs) from 3 to 5 distinct experiments, including 12 biological repetitions. The data presented here support the data presented in Fig. 5.

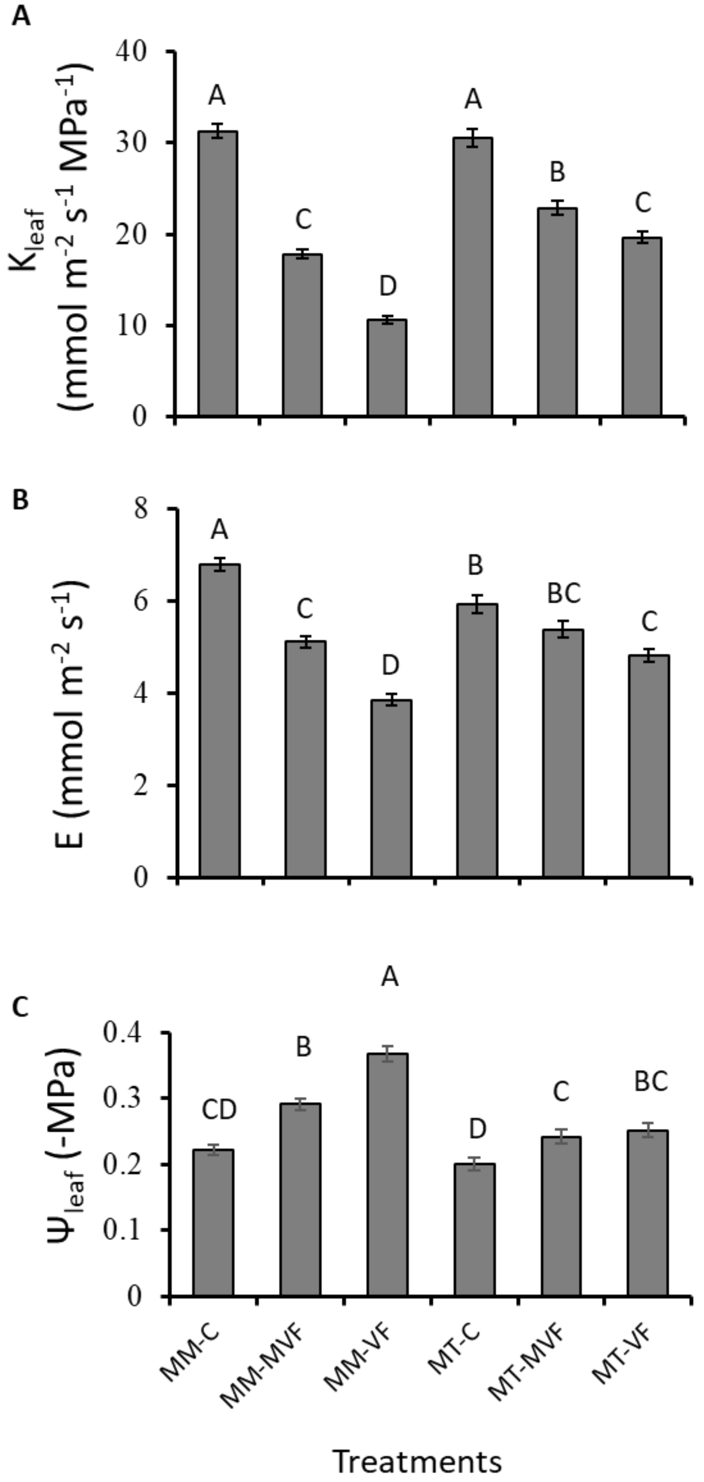


**Fig. S8 Effects of toxins released from the moderately virulent and virulent *F.oxysporum f. sp. lycopersici* strains on leaves detached from MM and MT tomato plants.**

A, Leaf hydraulic conductance, $K_{leaf}$, B, transpiration rate (E) and C, leaf water potential, $\Psi_{leaf}$ after 2–4 h of the xylem-fed toxin treatment. A, is identical to Fig. 5. C: Control groups fed with AXS alone. MVF and VF: Treated with toxins from the moderately virulent and virulent *F.oxysporum f. sp. lycopersici* strains, respectively. Different letters indicate significant differences between treatment, according to the Tukey-Kramer HSD test ($P < 0.05$). Data points are means (± SE) from 3 to 5 distinct experiments, with 29–30 biological repetitions overall. The data presented here support the data presented in Fig. 5.

**Table S1 Summary of experiments with different *F.oxysporum f. sp. lycopersici* strains and plants of varying levels of susceptibility.**

Days in the "transpiration", "mass" and "morphology" columns refer to the number of days after inoculation at which differences were first observed between inoculated and control plants.

Independent repetitions of each fungus × plant pairing are labeled with the same color. Plants tested: M82 – tolerant, R13 – susceptible, Mv – susceptible and MM – susceptible. Plants were inoculated with either the moderately virulent strain (mvF) or the highly virulent strain (vF). “Plant properties” are at day of infection. “Environmental properties” are the average for first 3 days after infection.

*Point in time (days from infection) at which a *t*-test indicated a significant difference relative to the control, $P < 0.05$

**Point in time (days from infection) at which 50% of plants exhibited chlorosis or wilting.

**Model S1 Logistic regression model of symptom occurrence.**

This logistic regression model predicts the likelihood of the occurrence of symptoms in plants inoculated with *F.oxysporum f. sp. lycopersici*. The model calculates the probability ('Prob[yes]_1' and 'Prob[no]_1') of symptom presence ('yes') or absence ('no') based on the daily light integral (DLI), plant average initial mass, relative humidity on the day of infection (RH day) and the specific combination of tomato cultivar ('Tomato') and Fusarium strain ('Fusarium'). Interaction effects between tomato cultivar and Fusarium strain are accounted for with differential coefficients. The final output, 'Most Likely Symptoms_1', determines the most probable symptom status based on the higher probability between 'yes' and 'no'. Coefficients are derived from the logistic regression equation ('Lin[yes]_1'), where the significance of each variable has been tested. Please note that this model is based on the data collected from the PlantArray system and is tailored to the conditions and variables specific to this study.

```
mp_1 = New Namespace("Fit Nominal Logistic - symthoms");
mp_1:predict = Function({DLI, Fusarium, Plant Av initial mass, RH day, Tomato},
  {Default Local},
  "Lin[yes]_1"n = (-2620.17699794658) + 58.2433796413645 * DLI + -52.6443864441494 *
Plant Av initial mass + 34.109539243059 * RH day
+Match( Fusarium,
        "mvF",
                Match( Tomato,
                        "M82", -92.261957864593,
                        "MM", 21.276227946165,
                        "Mv", 14.8915207305991,
                        "R13", 56.0942091878289,
                        .
                ),
        "vF",
                Match( Tomato,
                        "M82", 92.261957864593,
                        "MM", -21.276227946165,
                        "Mv", -14.8915207305991,
                        "R13", -56.0942091878289,
                ),      .
);
  "Prob[yes]_1"n = 1 / (1 + Exp( -"Lin[yes]_1"n ));
  "Prob[no]_1"n = 1 / (1 + Exp( "Lin[yes]_1"n ));
  Most Likely symthoms_1 = IfMax( "Prob[yes]_1"n, "yes", "Prob[no]_1"n, "no", "" );
);
```
